# Th1/17 Cells Infiltrate Murine Cytomegalovirus-Infected Renal Allografts via Virus-Induced CCL20 and Promote Th1 Cells through IL-17A

**DOI:** 10.1101/2021.08.05.455061

**Authors:** Ravi Dhital, Shashi Anand, Qiang Zeng, Victoria M. Velazquez, Srinivasa R. Boddeda, James R. Fitch, Ranjana W. Minz, Mukut Minz, Ashish Sharma, Rachel Cianciolo, Masako Shimamura

## Abstract

Cytomegalovirus (CMV) infection is associated with renal allograft failure by unknown mechanisms. In a murine renal transplant model, murine CMV (MCMV) induces intragraft infiltration of Th17 cells co-expressing Th1 cytokines, IFN-γ and TNF-α, but only a minority of intragraft Th17 cells are specific for MCMV antigens. Instead, MCMV promotes viral antigen-independent Th17 cell recruitment via CCL20-CCR6 and CXCL10-CXCR3 interactions. Th17 cells correlate directly with Th1 cell frequencies and inversely with Tregs in MCMV infected grafts. Pharmacologic inhibition of IL-17A reduces intragraft Th17 cells and neutrophils, increases Tregs, and reduces total but not MCMV-specific Th1 cells. Among a clinical renal transplant cohort with acute rejection, patients with CMV DNAemia had significantly higher serum IL-17A quantities compared to those without CMV DNAemia. Together, these findings indicate that CMV infection upregulates Th17 cell activity during acute rejection and suggests that inhibition of IL-17A may ameliorate CMV-associated allograft injury without impairing antiviral Th1 cells.

## INTRODUCTION

Human cytomegalovirus (HCMV) is a ubiquitous virus with seroprevalence up to 98% for adult populations worldwide^1, 2, 3^. Among healthy individuals, HCMV establishes latency with intermittent asymptomatic reactivations that are controlled by antiviral T cell responses^4, 5, 6^. However, HCMV reactivation can cause infectious complications in immunocompromised patients, such as solid organ transplant recipients. After organ transplantation, HCMV can cause either “direct effects” of viral disease (pneumonitis, hepatitis, colitis, retinitis, encephalitis) or “indirect effects” including increased risk for rejection, opportunistic infections, atherosclerosis, and post-transplant diabetes^7, 8, 9, 10^. CMV-specific T helper 1 (Th1) cells and cytotoxic CD8^+^ T cells are necessary for control of CMV infections, as shown in transplant patients and animal models utilizing murine CMV (MCMV) infection^11, 12, 13 14, 15, 16, 17^. The “direct effects” of HCMV are treated with antivirals such as valganciclovir, but antivirals cannot be used for long-term prophylaxis due to dose-limiting toxicities^18^. Short-term valganciclovir prophylaxis after renal transplantation is associated with lower graft loss rates compared to patients treated pre-emptively after onset of HCMV DNAemia, suggesting that viral reactivation contributes to graft injury^19^. Overall, however, the pathogenesis underlying HCMV “indirect effects” is not well understood.

In rodent renal transplant models, rat CMV and MCMV infection exacerbate allograft inflammation and accelerate fibrosis^20, 21, 22^. Similar to clinical observations, valganciclovir prophylaxis in murine transplants inhibits viral reactivation and improves late allograft fibrosis^23, 24^. Recently, our group showed that T helper 17 (Th17) cells are induced in MCMV-infected allografts, and that IL-6 inhibition reduces Th17 cell infiltrates and histopathologic allograft injury^22^. Th17 cells produce IL-17A/F and other proinflammatory cytokines including IL-21, IL-22, and TNF-α, which contribute to inflammatory responses during autoimmune and infectious diseases and are also observed during allograft rejection^25, 26, 27, 28, 29^. Th17 cells, IL-17A cytokine quantities and mRNA transcripts are elevated in allografts and blood of kidney and liver transplant patients with acute rejection, as are urinary levels of the Th17 cell chemoattractant, CCL20 ^30, 31, 32, 33 34, 35, 36, 37, 38^. In a murine cardiac transplant model, IL-17 mRNA transcripts, IL-17 cytokine levels and Th17 cell frequencies are also elevated in acutely rejecting grafts^39, 40^. However, these studies did not analyze the contribution of CMV to Th17 cells during acute rejection.

Th17 cells have known effector functions against extracellular pathogens, mucoepithelial infections, and intracellular bacteria, but are not typically associated with antiviral immunity^41, 42, 43^. However, Th17 cells have been reported to contribute to pathological inflammation in experimental and clinical viral infections, including influenza pneumonia and liver infection with hepatitis B and C viruses^44, 45, 46^. We therefore postulated that Th17 cells might contribute to pathological inflammation during acute rejection of MCMV infected renal allografts. To examine this, we investigated the phenotype of MCMV-induced Th17 cells for viral antigen specificity, intragraft conditions promoting their generation, and Th17 cell effector cytokine activity within rejecting murine renal allografts. The association of HCMV DNAemia with serum IL-17A detection was assessed among a cohort of renal transplant patients with acute rejection.

## RESULTS

### MCMV induces graft-infiltrating Th17 cells co-expressing Th1 cytokines

In our prior studies, murine cytomegalovirus (MCMV) induced the infiltration of Th17 cells into murine renal allografts at day 14 post-transplantation^22^. To examine earlier events contributing to Th17 cell infiltration, allogeneic renal transplants were performed using donor (D) and recipient (R) MCMV negative (D-R-) and positive (D+R+) combinations with cyclosporine immunosuppression, sacrificed at day 7 post-transplant (Figure 1A), and frequencies of graft-infiltrating CD4^+^ T- cells expressing IL-17A, IFN-γ and TNF-α were quantified by intracellular cytokine staining/flow cytometry (ICS/FC) following PMA/ionomycin stimulation (Figures 1B and S1). Consistent with the findings at day 14, D+R+ allografts at day 7 had significantly higher frequencies of total IL-17A^+^ Th17 cells compared to D-R-allografts, as well as higher frequencies of Th17 cells co-expressing Th1 cytokines, IFN-γ and/or TNF-α (Figure 1C). D+R+ grafts also had higher concentrations of IL-17A and IFN-γ than D-R-grafts, whereas TNF-α quantities were similar (Figure 1D). Comparing IL-17A expressing CD4^+^, CD8^+^ and MHCII^+^ cells in D+R+ and D-R-allografts, only Th17 cells were more abundant in D+R+ allografts (Figure 1E), indicating that Th17 cells are the most likely source of IL-17A observed in D+R+ allografts. Compared to spleens, D+R+ allografts had significantly higher frequencies of Th17 cells expressing IL-17A alone or in combination with Th1 cytokines (Figure 1F), indicating that the phenotype of graft-infiltrating Th17 cells differs from that observed systemically. Similarly, allografts had significantly higher quantities of IL-17A, IFN-γ and TNF-α cytokines compared to spleens. Together, these results indicate that MCMV infection induces the infiltration of Th17 cells with a Th1/17 profile, which are associated with higher intragraft quantities of IL-17A and IFN-γ.

**Figure 1.**
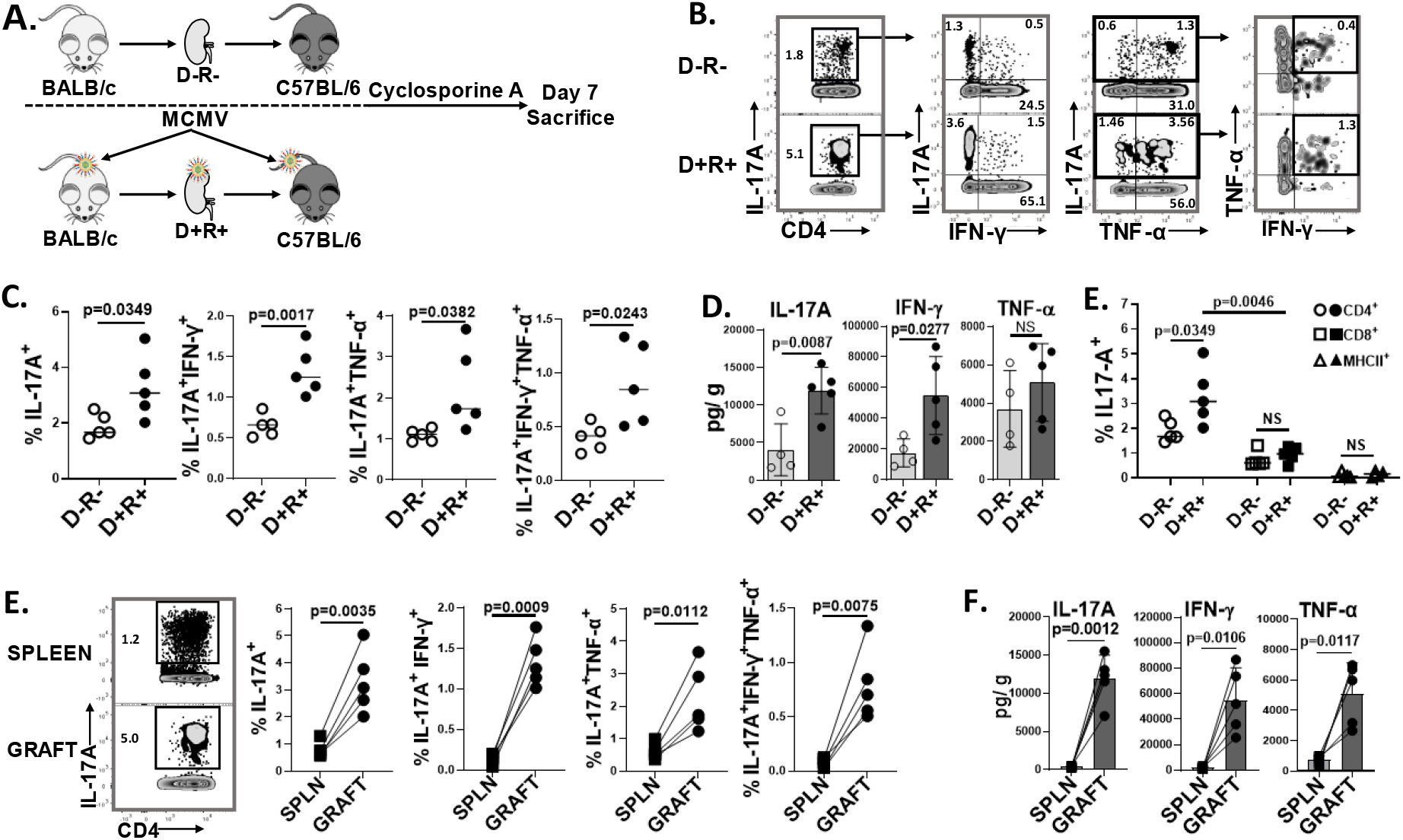
MCMV induces intragraft infiltration of Th17 cells co-expressing Th1 cytokines. (A) Study design. Allogeneic renal transplantation was performed between BALB/c and C57BL/6 mice with cyclosporine immunosuppression. On day 7, splenocytes and intragraft leucocytes were analyzed by flow cytometry. (B) Representative flow cytometry plots showing frequencies of CD4^+^IL-17A^+^ (Th17) cells and cytokine profiles. (C) Frequencies of cytokine-expressing Th17 cells were compared for D-R- and D+R+ allografts. (D) Intragraft quantities of IL-17A, IFN-γ and TNF-α (picogram per gram tissue, pg/g) in D-R- and D+R+ allografts. (E) Frequencies of IL-17A expressing CD4^+^, CD8^+^ and MHC-II^+^ leucocytes in D-R- and D+R+ allografts. (F) Representative flowplots and frequencies of Th17 cells in D+R+ spleens (SPLN) and allografts (GRAFT). (G) Cytokine quantities (pg/g) in D+R+ spleens and allografts. All data are represented as mean ± standard deviation (SD) and analyzed by two- sided Student’s t-test. NS, not significant (p>0.05).

### MCMV antigen-specific Th17 cells differ phenotypically from viral antigen-independent Th17 cells

To determine whether Th17 cells in D+R+ allografts have specificity for MCMV antigens, we first compared detection of MCMV-specific Th17 cells among D+R+ splenocytes after stimulation with either viral lysate or a pool of class II (I-A^b^)-restricted MCMV peptides (Figure S2A and Table S3) ^47^. As MCMV-specific Th1 and Th17 cells could be detected at comparable or higher frequencies after peptide stimulation (Figure S2B), the peptide pool was used for subsequent experiments.

The Kinetics of MCMV-specific Th17 cells are hitherto uncharacterized. Therefore, we first used non-transplant B6 mice to analyze the anti-MCMV Th17 immune response after primary infection. B6 mice were infected with MCMV at 1 × 10^6^ plaque-forming units (pfu), and splenocytes isolated at days 0, 7, 14, 21, and 28 were incubated with PMA or MCMV peptide pool and analyzed for IL-17A expression (Figure 2A). MCMV-specific Th17 cells increased at day 7, comprising an average of 13.55 ± 4.67% of the total Th17 cells (Figure 2B), and rapidly returned to baseline levels from day 14 onward (Figure 2A). We next identified MCMV-specific graft-infiltrating Th17 cells from D+R+ transplants (Figures 2C-E). Unexpectedly, MCMV specific Th17 cells comprised only a small percentage (5.48 ± 0.97%) of total graft-infiltrating Th17 cells (Figure 2D) and predominantly expressed IL-17A alone or with IFN-γ, but not TNF-α (Figure 2E). As the majority of MCMV-induced Th17 cells lacked specificity for viral antigens, we next examined the requirement for any graft-specific antigens, by performing transplants using recipient OTII transgenic mice, which express the CD4 co-receptor specific for chicken ovalbumin (OVA_323-339_) in context of I-A^b^ (Figures 2F and S3). D- and D+ donor kidneys, which lack OVA expression, were analyzed at day 7 post-transplant for I-A^b^-OVA_323-339_ tetramer^+^ Th17 cells. OVA tetramer^+^ Th17 cells were significantly higher in the D+ grafts (Figure 2G), indicating that MCMV infection can induce infiltration of Th17 cells lacking specificity for any allograft-expressed antigens. Tetramer^+^ Th17 cells co-expressed TNF-α with or without IFN-γ, similar to the PMA+ but not MCMV peptide+ Th17 cells observed in wild-type D+R+ transplants (Figure 2H). Together, these results indicate that MCMV infection can induce infiltration of both viral antigen-dependent and antigen-independent Th17 cells that differ in their cytokine expression profiles.

**Figure 2.**
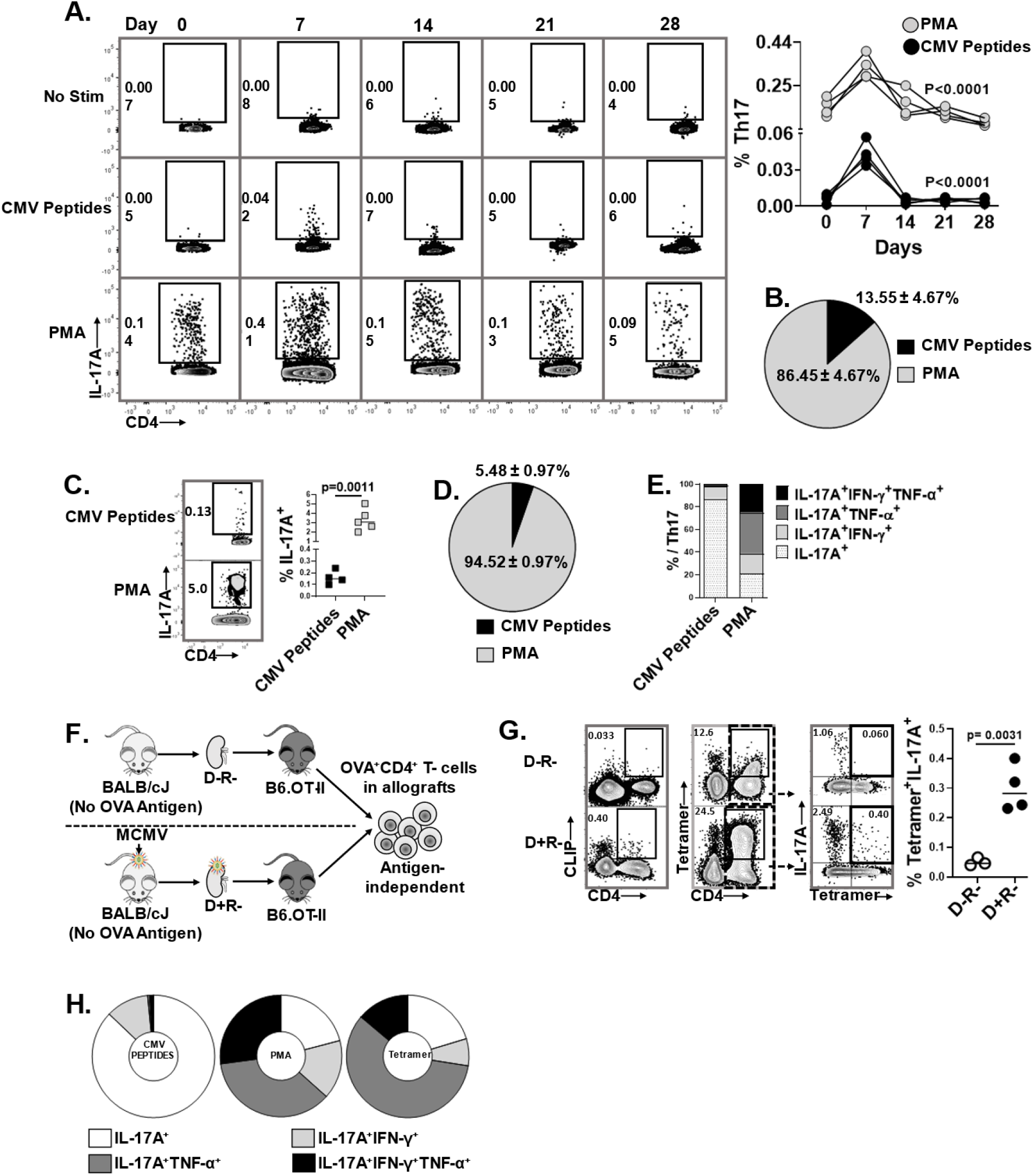
MCMV-specific and antigen-independent Th17 cells infiltrate virus infected allografts. (A, B) Non-transplant B6 mice were infected with MCMV on Day 0 and splenic Th17 cell frequencies were quantified at days 0, 7, 14, 21, and 28 post-infection. Splenocytes were stimulated with either PMA or MCMV peptides and stained for IL-17A expressing CD4^+^ T- cells. (A) Representative flow plots showing frequencies of CMV specific and total Th17 cells at indicated days; graph shows frequencies of total (gray circles) and MCMV-specific (black circles) Th17 cells over time. (B) Pie chart shows the percentage of MCMV-specific Th17 cells in non-transplant spleen at day 7 post-infection. (C) Representative flow plots and frequencies of CMV-specific (CMV peptides+) and total (PMA+) Th17 cells in allografts of D+R+ transplant recipients. (D) Pie chart shows percentage of MCMV-specific Th17 cells in allografts. (E) Proportions of intragraft Th17 cells expressing IL-17A, IFN-γ and/or TNF-α were compared for MCMV-specific and total Th17 cells. (F) Experimental design. B6.OT-II transgenic recipients received D- or D+ allografts lacking expression of OVA antigen, so that OVA^+^ Th17 cells are recruited to allografts by antigen-independent mechanisms. (G) OVA-specific Th17 cells were detected using I-A^b^-OVA_323-339_-APC tetramer staining, with human CLIP-APC tetramer used as control (Supplementary Fig. 3). Representative flow plots show tetramer staining of CD4^+^ T cells derived from D-R- and D+R+ allografts. Graph shows the frequencies of Tetramer^+^ Th17 cells compared between the groups. (H) Cytokine expression profiles were compared for intragraft MCMV specific and PMA+ Th17 cells in wild-type recipients, and for Tetramer^+^ Th17 cells from OTII recipients. Data are represented as mean ± standard deviation (SD) and are analyzed by two-sided Student’s t-test (C, G) or one-way ANOVA (A). PMA, Phorbol 12-myristate 13-acetate; CLIP, Class II-associated invariant chain peptide.

### The microenvironment of MCMV infected allografts favors Th17 cell recruitment

The presence of antigen-independent OVA-specific Th17 cells in allografts suggests that MCMV might induce expression of cytokines promoting Th17 cell infiltration. To define the microenvironment of transplantation, we first compared the gene expression profiles of D+ allografts to those of MCMV infected non-transplant BALB/c kidneys by RNAseq. Ingenuity pathway analysis (IPA) showed that transplantation of MCMV-infected kidneys led to significant changes in 5502 genes (adjusted p values <0.05), involved in 391 canonical pathways. The Th17 activation pathway showed 107 differentially expressed genes. Transplant kidneys showed higher expression of genes encoding transcription factors, cytokines, and cytokine receptors that are required for differentiation of naïve CD4^+^ T- cells to Th17 cells (Figure 3A), as well as transcripts for chemokines and chemokine receptors involved in the recruitment of Th17 cells, including *CXCL10, CCL20, CXCR3, and CCR6* (Figure 3B). Similar findings were identified in comparisons of non-transplant uninfected kidneys and D- allografts (Figure S4). D- and D+ allografts had no significantly differentially expressed Th17 pathway associated transcripts, nor were differences observed between non-transplant MCMV uninfected and infected kidneys. These analyses indicate that the D+ and D- allograft microenvironment is favorable for Th17 development and/or recruitment compared to native kidneys.

**Figure 3.**
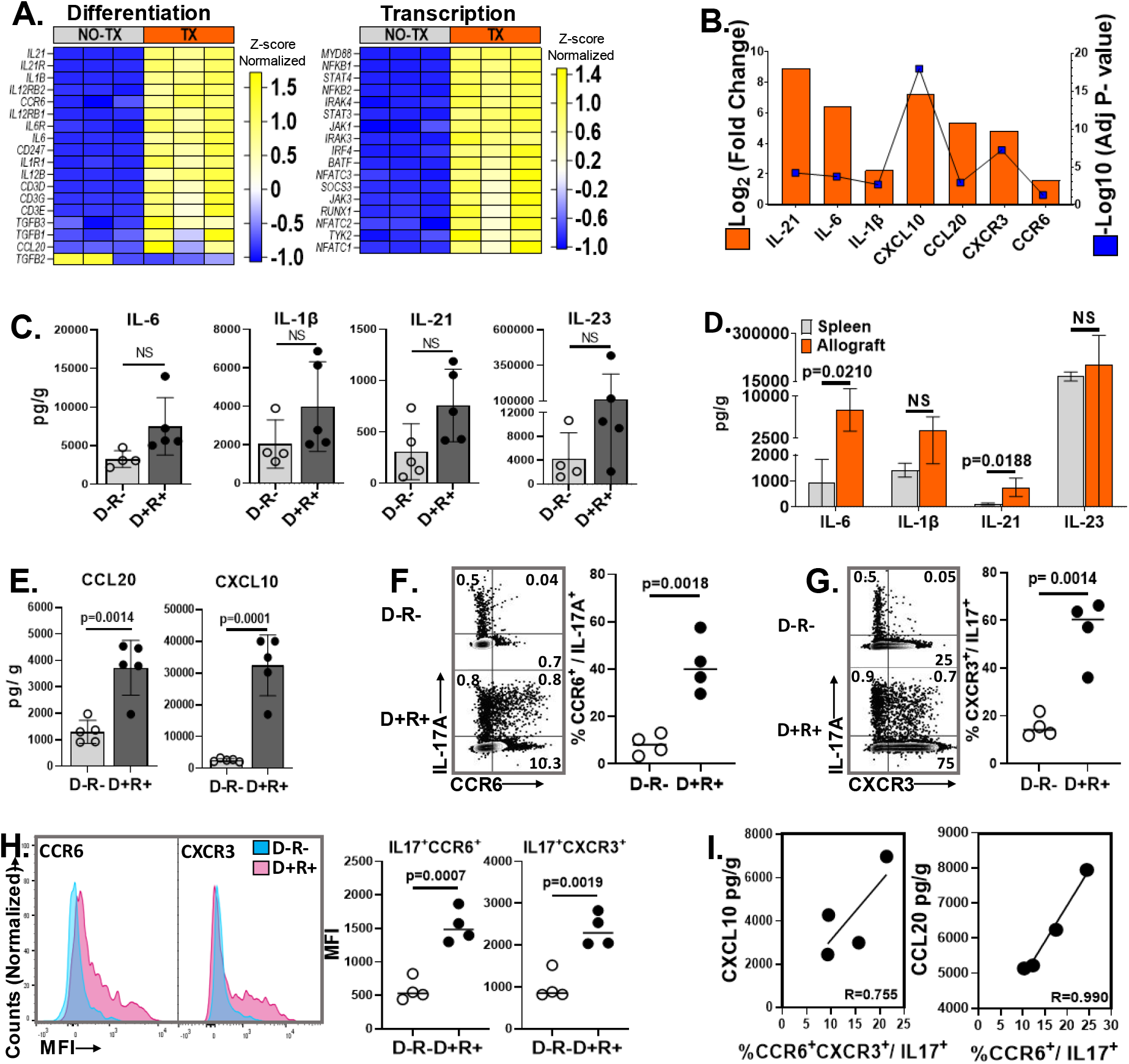
MCMV infected allograft microenvironment favors Th17 cell recruitment. (A, B) Allografts from D+ transplants (TX) and MCMV infected BALB/c kidneys (NO-TX) were compared for gene expression by RNA-seq. (A) Heat map shows differentially expressed genes encoding molecules required for Th17 differentiation and transcription factors. Each column represents a single sample whereas the rows represent intensities of gene expression. (B) Transcripts for Th17 cell differentiating cytokines and recruiting chemokines are upregulated in D+ transplants. (C) Comparison of intragraft Th17 differentiating cytokine quantities between D-R- and D+R+ transplants. (D) Quantities of Th17 cell differentiating cytokines in D+R+ spleens and allografts. (E) Comparison of Th17 cell recruiting chemokine quantities in D-R- and D+R+ allografts. (F, G) Representative flow plots and frequencies of CCR6^+^ and CXCR3^+^ Th17 cells in D-R- and D+R+ allografts. (H) Mean fluorescence intensity (MFI) of CCR6 and CXCR3 expression for Th17 cells from D-R- (Blue) and D+R+ (Pink) allografts. (I) Correlation between intragraft chemokines and receptors expressed by Th17 cells from D+R+ transplants. All data are represented as mean ± standard deviation (SD) and are analyzed by two- sided Student’s t-test or Pearson correlation. NS, not significant (p>0.05).

Having confirmed that transplantation induces upregulation of numerous Th17 cell-associated transcripts, we next compared the phenotypic expression of Th17 cell-associated cytokines and chemokines in D-R- and D+R+ allografts. The Th17 cell differentiating cytokines, IL-6 and IL-21, trended higher in D+R+ allografts as compared to D-R- allografts (p=0.0664 and p=0.0538, respectively), but none reached statistical significance (Figure 3C), whereas IL-6 and IL-21 were significantly higher in D+R+ allografts than spleens (Figure 3D). However, D+R+ allografts had significantly higher quantities of Th17 cell-recruiting chemokines, CCL20 and CXCL10, compared to D-R- allografts (Figure 3E). Th17 cells in D+R+ allografts expressed CCR6 and CXCR3 at both, higher frequencies and higher MFIs compared to Th17 cells from D-R- allografts (Figures 3F-H and S5A). Frequencies of CCR6^+^CXCR3^+^ and CCR6^+^ Th17 cells correlated with quantities of their respective ligands, CXCL10 and CCL20 (Figure 3I). Together, these findings indicate that the chemokine microenvironment of MCMV-infected allografts favors recruitment of CCR6^+^ and CCR6^+^CXCR3^+^ Th17 cells via expression of CCL20 and CXCL10.

### Ischemia Reperfusion injury (IRI), MCMV infection, and allogeneic transplantation each contribute to Th17 cell recruitment

Ischemia-reperfusion injury (IRI) alone can generate trafficking signals for CD4^+^ T cells into allografts and can also induce MCMV reactivation from latently infected transplant organs^23, 48, 49^. We therefore interrogated the contribution of IRI without alloimmune responses by performing syngeneic transplants using D-R- (IRI alone) and D+R+ (IRI + MCMV) kidneys (Figure 4A). Both CCL20 and CXCL10 were detected in syngeneic D-R- transplants (Figure 4B), as well as Th17 cell infiltrates (Figure 4C), confirming that IRI alone can induce Th17 cell recruitment. Syngeneic D+R+ grafts had higher quantities of CCL20, CXCL10, and Th17 cells compared to syngeneic D-R- grafts (Figures 4B-C), indicating that MCMV reactivation after IRI increases Th17 cell recruitment even in the absence of allogeneic signals. Direct comparison of Th17 cell frequencies and cytokine-expressing subsets (Figure 4D) shows that IRI, MCMV infection, and allogeneic transplantation each contribute to Th17 cell recruitment but is greatest after allogeneic transplantation of MCMV infected donor organs (Figure 4E).

**Figure 4.**
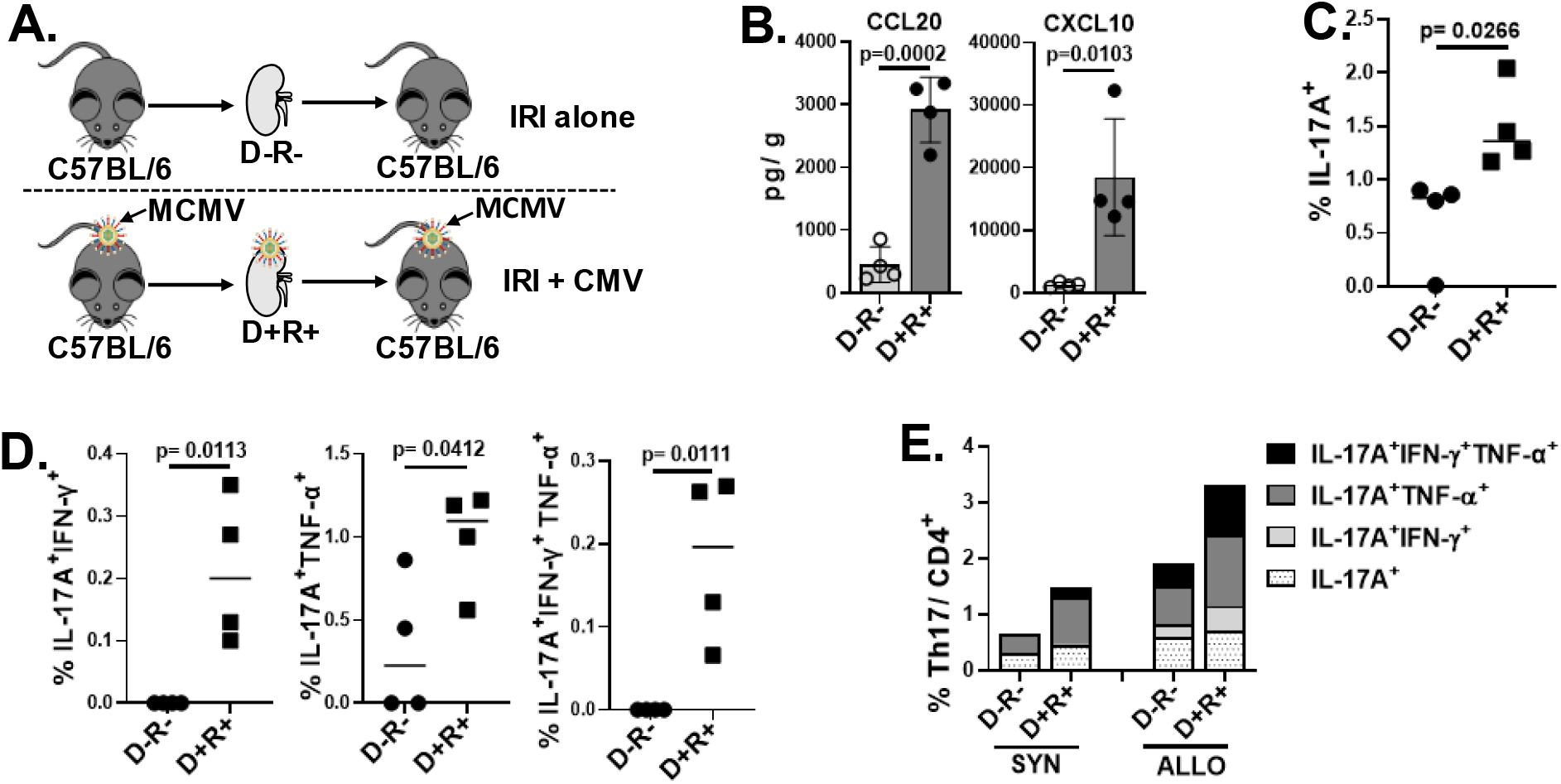
Ischemia Reperfusion Injury, MCMV infection, and allogeneic transplantation each contribute to Th17 cell infiltration into allografts. (A) Syngeneic transplantation was performed using D-R- and D+R+ grafts to evaluate role of ischemia-reperfusion injury (IRI) without alloimmune responses. (B) CCL20 and CXCL10 expression in D-R- and D+R+ syngeneic transplants. (C) Frequencies of Th17 infiltrates in syngeneic D-R- and D+R+ grafts. (D) Intragraft Th17 cells co-expressing IFN-γ and/or TNF-α in D-R- and D+R+ grafts. (E) Percentage of single or multiple cytokines expressing Th17 cells in syngeneic and allogeneic grafts. Except that only mean values are shown in figure E, all data are represented as mean ± standard deviation (SD) and are analyzed by two- sided Student’s t-test.

### Th17 cells are associated with graft-infiltrating Th1 cells and reduced Tregs

Signals promoting development of Th17 cells can suppress T helper 1 (Th1) and regulatory T (Treg) subsets^50, 51^. However, IPA analysis of RNA-seq data revealed that the Th1/Th2 activation pathway was the second most highly upregulated pathway in the list of total 391 enriched canonical pathways (Table S4), consistent with studies associating Th1 cells with acute T cell mediated rejection in clinical transplantation^52, 53, 54^. D+R+ allografts showed higher expression of 64 genes encoding proteins and transcription factors involved in the Th1 pathway, including *IFN-γ* (Figure 5A). D+R+ allografts had higher frequencies of Th1 cells compared to D-R- allografts (Figure 5B) and D+R+ spleens (Figure 5C), and intragraft Th1 cell frequencies correlated with Th17 cell frequencies (Figure 5B). The Th17:Th1 ratio was higher in D+R+ allografts compared to spleens (Figure 5D), indicating that the proportion of Th17 to Th1 cells was relatively higher in allografts. Comparing frequencies of Foxp3 and interleukin-10 (IL-10) expressing CD4^+^ T cells (Tregs) in allografts and spleens (Figure S1), D-R- allografts had higher frequencies of Tregs compared to D+R+ grafts, resulting in a higher Th17: Treg ratio in D+R+ allografts (Figures 5E-F). Similar comparisons between D-R- and D+R+ spleens (Figures 5G-H) showed a higher frequency of Foxp3^+^ (Figure 5G) and IL-10^+^ (Figure 5H) Tregs in D-R- spleens corresponding with a lower Th17: Treg ratio (Figure 5H). Taken together, these results demonstrate that Th17 cells do not inhibit Th1 cells in this model but rather correlate directly with Th1 cell infiltrates, whereas Treg activity is reduced relative to Th17 cells.

**Figure 5.**
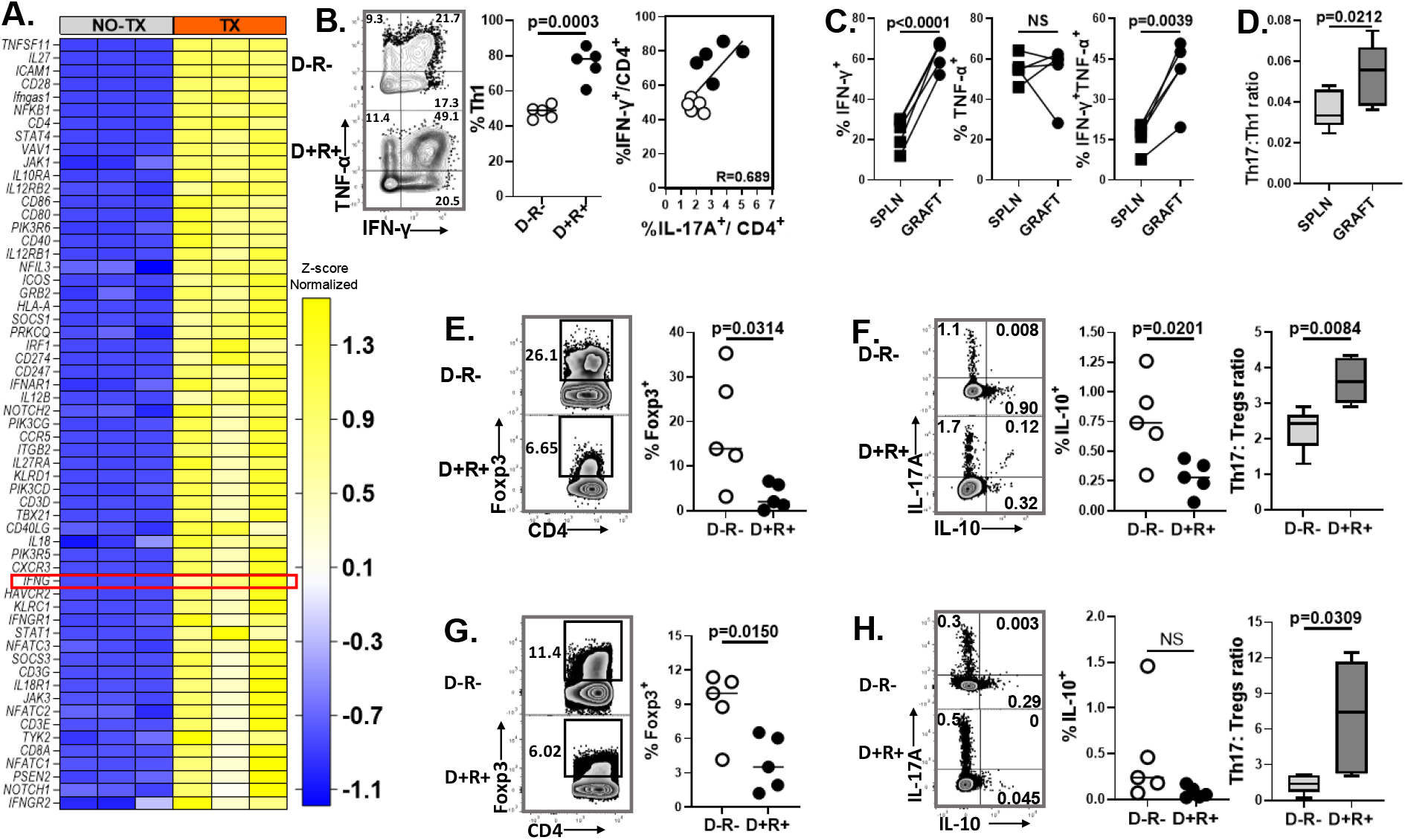
MCMV infection is associated with increased Th1 cells and decreased Tregs. (A) Heat map shows differential expression of transcripts involved with Th1 cell activation in D+ allografts (TX) compared to non-transplant MCMV infected kidneys (NO TX). (B) Representative flow plots and frequencies of Th1 cells in D-R- and D+R+ allografts. Correlation graph shows positive correlation between intragraft frequencies of Th1 and Th17 cells in D+R+ transplants. (C) Frequencies of IFN-γ and/or TNF-α expressing Th1 cells in D+R+ spleens and allografts. (D) Ratio of Th17:Th1 cell infiltrates in D+R+ spleens and allografts. (E-H) Representative flow plots and frequencies of Foxp3^+^ Tregs and IL-10 expressing Tregs, and box plots shows ratio of Th17: Treg cells in allografts (E, F) and spleens (G, H). All data are represented as mean ± standard deviation (SD) and are analyzed by two- sided Student’s t-test or Pearson correlation. NS, not significant (p>0.05).

### Inhibition of IL-17A reduces infiltration of neutrophils and Th1 cells, promotes Tregs, and ameliorates allograft injury

IL-17A mobilizes neutrophils to sites of inflammation through induction of chemokines, CXCL1 and CXCL5^55, 56, 57^. Comparing D-R- and D+R+ allografts, CXCL1 was significantly higher in D+R+ allografts (Figure 6A) and correlated with intragraft Th17 cell frequencies (Figure 6B), but CXCL5 was not significantly different (Figure 6A). These results suggested that IL-17A might promote neutrophil recruitment to MCMV infected allografts. To confirm this, a cohort of allogeneic D+R+ recipients were treated with either anti-IL-17A antibodies or isotype control antibodies (Figure 6C). Anti-IL-17A treated animals had significantly reduced frequencies of Th17 cells in allografts, spleen, and peripheral circulation compared to isotype treated mice (Figure 6D), as well as reduced CD11b^+^Ly6G^+^ neutrophil populations in allografts, but not in spleens and peripheral blood (Figures 6E and S5B). Anti-IL-17A treated grafts also had significantly increased Foxp3^+^ Treg frequencies (Figure 6F), with decreased Th17: Tregs ratio (Figure 6G). Unexpectedly, Th1 cell infiltrates also were lower in anti-IL-17A treated allografts compared to control grafts (Figure 6H), resulting in a decreased Th1: Treg ratio in the treatment group (Figure 6I). The anti-IL-17A treatment selectively decreased total but not MCMV antigen-specific Th1 cells within allografts (Figure 6J) and had no effect upon splenic Th1 cells (Figure 6K). Anti-IL17A treated allografts also had less severe glomerulitis, tubulitis, and arteritis by histopathology (Damage Score, 17.5 ± 2.1) compared to isotype treated grafts (Damage Score, 20.0 ± 1.4) at day 14 post-transplantation (Figure 6L).

**Figure 6.**
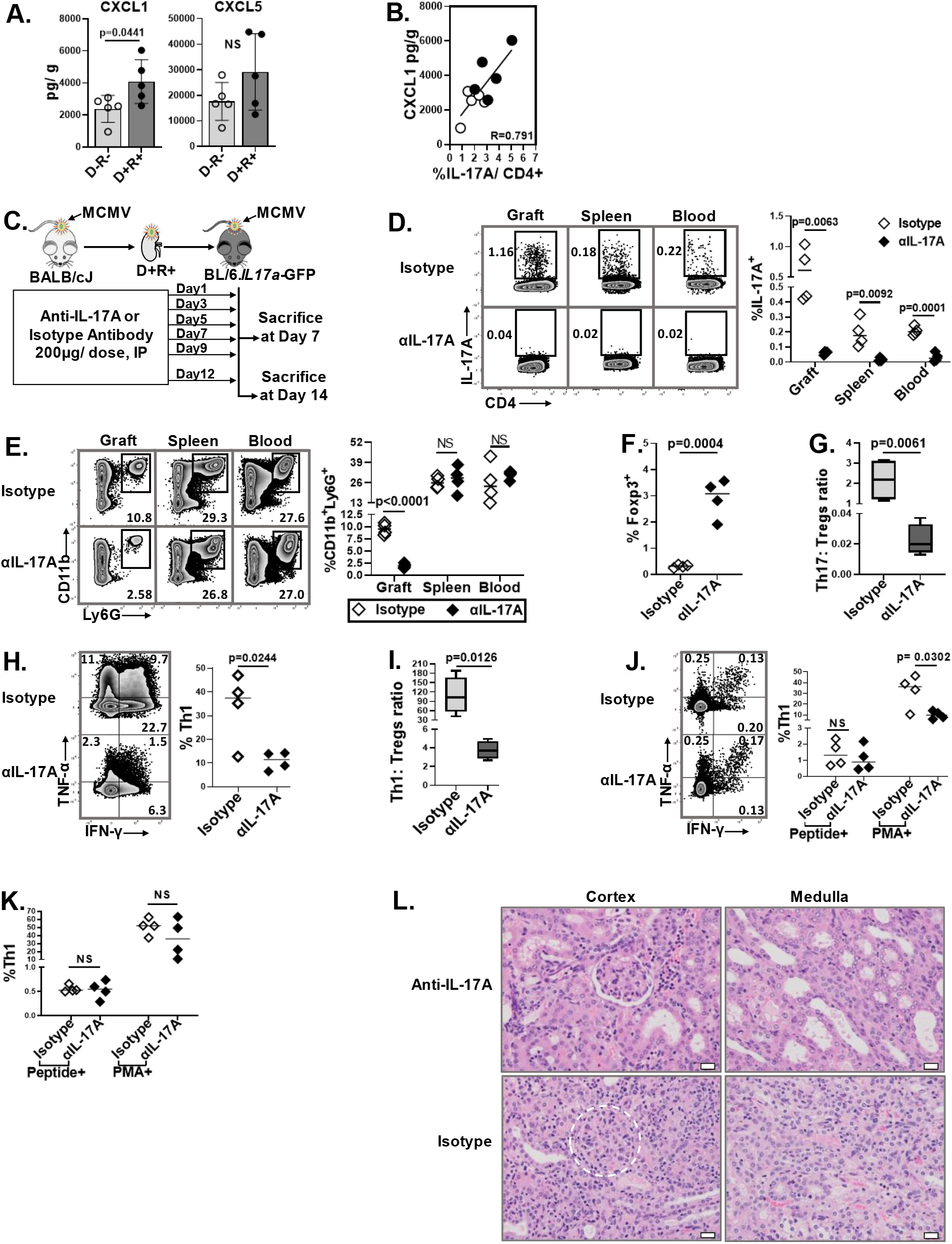
IL-17A inhibition modulates neutrophil and T helper cell infiltrates. (A) IL-17A induced neutrophil chemoattractants, CXCL1 and CXCL5, were compared for D-R- and D+R+ allografts. (B) Correlation between intragraft CXCL1 and Th17 infiltrates. (C) Study design. D+R+ recipients were treated with anti-IL-17A antibodies or isotype control antibodies at 200µg/dose on indicated days post-transplantation and sacrificed at day 7 or 14. (D-K) Day 7 post-transplantation. (D) Representative flow plots and frequencies of Th17 cells in allograft, spleen, and peripheral blood of recipient mice from anti-IL-17A treated (αIL-17A) and isotype control groups. (E) Representative flow plots and frequencies of CD11b^+^Ly6G^+^ neutrophils in organs and blood of anti-IL-17A and isotype treated transplant recipients. (F) Frequencies of Foxp3^+^ Tregs in allografts of anti-IL17A treated and control mice. (G) Intragraft Th17: Tregs ratio between the groups. (H) Representative flow plots and frequencies of intragraft Th1 cell infiltrates between the groups. (I) Ratio of Th1: Tregs in allografts. (J, K) Representative flow plots and frequencies of MCMV-specific and -nonspecific Th1 cells in allografts (J) and spleens (K). (L) Hematoxylin and Eosin (H&E) staining of allografts from anti-IL-17A treated and control groups (40x). White dotted circle, glomerulus. White bar, 20µm. All data are represented as mean ± standard deviation (SD) and are analyzed by two- sided Student’s t-test or Pearson correlation. NS, not significant (p>0.05).

### HCMV DNAemia is associated with elevated serum IL-17A quantities during acute rejection in clinical renal transplantation

To extend observations in the murine model to clinical renal transplantation, we retrospectively analyzed samples collected prospectively from a cohort of renal transplant patients during the first-year post-transplant at a tertiary referral hospital in India (Figure S6). Of 211 recruited recipients, 35 recipients had biopsy proven acute allograft rejection (AR) in the first year of transplantation, whereas the remainder (N=173) had no acute rejection (NR). AR was classified histopathologically according to Banff criteria as acute cellular rejection (N= 20), acute antibody mediated rejection (N= 6) and acute mixed rejection (N= 9). AR patients with a first episode of acute cellular or mixed rejection, who had blood samples with sufficient RNA quality for analysis (N=24) were matched with NR patients (N=29) by age and sex. The demographics of the AR and NR groups are shown in Table S1. The mean time for onset of acute rejection (AR) was 32.33 ± 49.46 days post-transplant. Samples of NR patients were compared at the post-transplant timepoint corresponding to the timepoint at which AR occurred in the matched case patient. Recipients with AR had significantly higher serum IL-17A quantities at the time of rejection compared to NR recipients (Figure 7A), which remained significantly elevated through the subsequent 2 timepoints after the AR event. Quantitative reverse-transcriptase PCR of whole blood showed higher RORyt and lower FOXP3 mRNA among AR patients during rejection episodes (Figures 7B, D and F). Conversely, serum IL-10 (Figure 7C) was lower among AR patients than NR patients, reflected as higher IL-17A:IL-10 ratio at 1- and 3-months post-transplantation (Figure 7E). Together, these findings are consistent with published reports describing Th17 cell activity during clinical AR ^34^.

**Figure 7.**
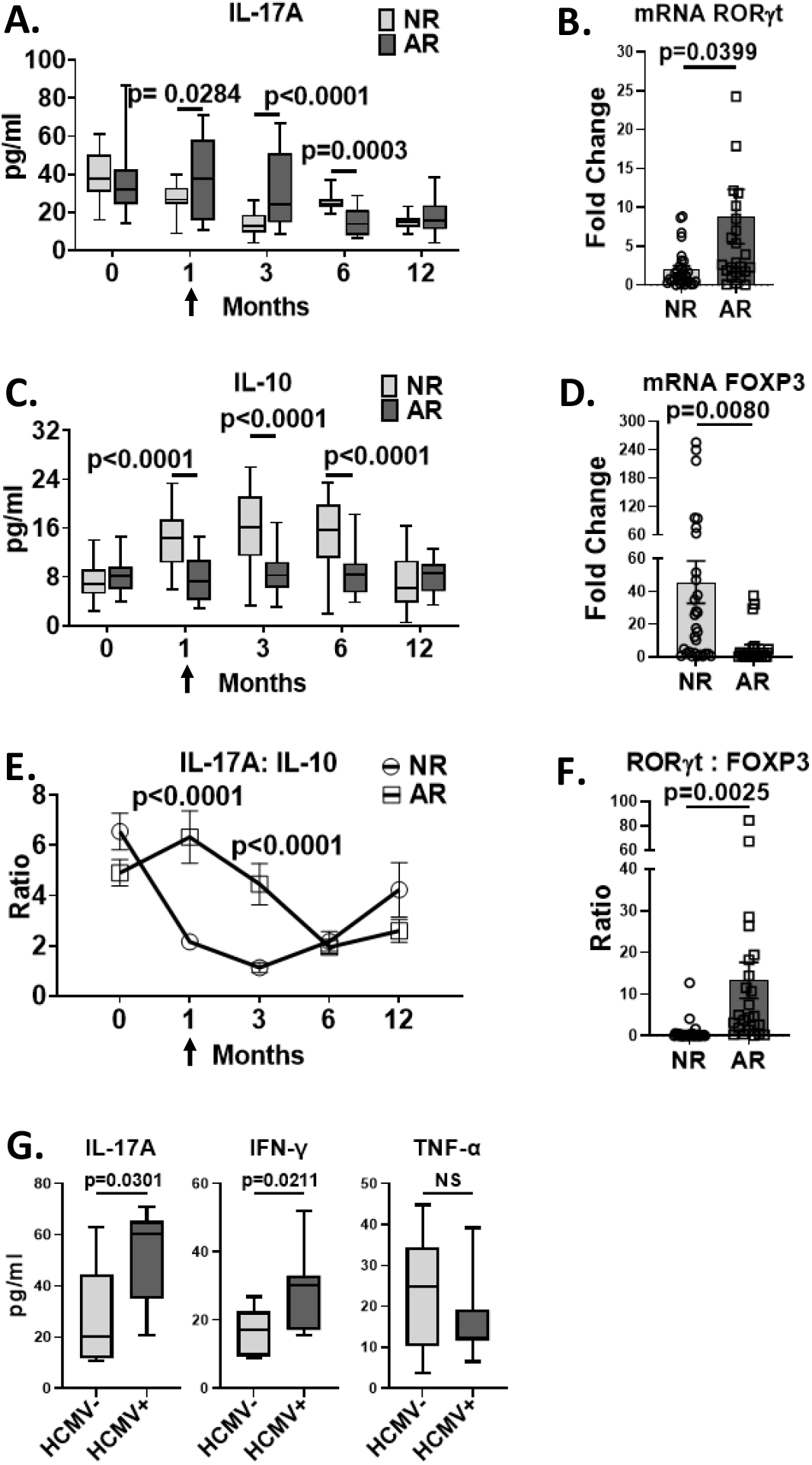
CMV DNAemia is associated with elevated serum IL-17A and IFN-γ in kidney trans lant atients with acute rejection. Clinical renal transplant recipients with acute rejection (AR, N=24) were compared to those with no rejection (NR, N=29) over the first 12 months post-transplant. Blood was analyzed at pre-transplant (0), 1-, 3-, 6- and 12-months post-transplant and at the time of rejection (arrow). (A, C, E) Serum cytokine levels were measured at indicated post-transplant timepoints. (B, D) Expression of transcription factors were quantified in whole blood by RT-PCR. (F) Ratio of mRNA expression of RORγt and Foxp3 were compared at the time of rejection for AR and NR groups. (G) HCMV DNAemia in the AR group was determined by quantitative DNA PCR. Serum cytokine quantities were compared for patients with (HCMV+) and without (HCMV-) HCMV DNAemia. All data are represented as mean ± SEM and are analyzed by two- sided Student’s t-test. NS, not significant (p>0.05).

We next examined HCMV DNAemia among 20 AR patients and 17 NR patients for whom serum was available for HCMV DNA PCR testing. In the AR group, 40% (8/20) had HCMV DNAemia at the time of AR, compared to 6% (1/17) of the NR patients (p= 0.0159) (Table S2), consistent with HCMV reactivation during AR. Within the AR group, serum IL-17A and IFN-γ were significantly higher among patients with HCMV DNAemia (HCMV+) compared to those without HCMV DNAemia (HCMV-), but no difference was observed for TNF-α (Figure 7G). This cytokine profile strikingly resembles that observed in MCMV+ allografts in the murine model (Figure 1D) and supports that HCMV reactivation is associated with IL-17A and IFN-γ expression during acute rejection.

## DISCUSSION

We have previously shown that MCMV infected allografts have higher Th17 cell infiltrates compared to uninfected grafts, and that treatment with anti-IL-6 antibodies reduces Th17 cell infiltrates and ameliorates MCMV-induced histopathologic damage^22^. In this study, the phenotype, antigen specificity, and microenvironmental cues associated with MCMV induced, graft infiltrating Th17 cells were characterized. Intragraft MCMV-induced Th17 cells co-express Th1 cytokines that differ according to antigen specificity, with MCMV-specific Th17 cells co-expressing IFN-γ but not TNF-α, and viral antigen-independent Th17 cells co-expressing TNF-α with or without IFN-γ. Unexpectedly, MCMV-specific Th17 cells comprise only a small minority of total graft-infiltrating Th17 cells. Instead, the microenvironment of MCMV infected allografts favors recruitment of MCMV antigen-independent Th17 cells via CCL20-CCR6 and CXCL10-CXCR3 interactions. These microenvironmental cues are also observed in MCMV-infected syngeneic grafts, indicating that MCMV reactivation after ischemia-reperfusion injury (IRI) can facilitate Th17 cell recruitment in the absence of allogeneic stimuli. MCMV-induced Th17 cells are also associated with lower Tregs frequencies and higher Th1 cell infiltrates, promoting the pro-inflammatory milieu of acute rejection. Inhibition of the Th17 cell effector cytokine, IL-17A, reduces neutrophil recruitment, increases Treg frequencies, and reduces Th1 cell frequencies. Together, these findings indicate that MCMV infection exacerbates recruitment of Th17 cells to allografts not only via expression of viral antigens, but also by altering the chemokine microenvironment. Anti-IL17A treatment ameliorates MCMV-associated allograft injury by modulating T helper cell and neutrophil infiltrates.

Renal cells, such as endothelial, mesangial, and tubular epithelial cells (TECs) can secrete a wide range of chemokines such as monocyte chemoattractant protein 1 (MCP1/CCL2), interleukin (IL-8), CXCL10 (IP-10), RANTES (CCL5), neutrophil-activating protein-3 (NAP-3/CXCL1), macrophage inflammatory protein-1 alpha (MIP-1α/CCL3), MIP-2α (CXCL2) and MIP-3α (CCL20), under the influence of proinflammatory stimuli including TNF-α, IL-1β, IFN-γ, IgG and IgA, and ischemia-reperfusion injury^58, 59, 60, 61^. IRI-induced chemokines and cytokines such as IL-6, IL-1β, and TNF-α also promote CMV reactivation *in vitro* and after syngeneic and allogeneic transplantation^23, 49, 62, 63, 64, 65^. After viral reactivation, CMV infection induces expression of IL-6, IL-8, and TNF-α from monocytes, TGF-β1 from renal tubular epithelial cells, and IL-8, CXCL11, RANTES, IL-1β, IL-6, fractalkine (CX3CL1), and CXCL1 from endothelial cells *in vitro*^66, 67, 68, 69^. Rat CMV infection also increases expression of the chemokines RANTES, MCP-1, MIP-1α and IP-10/CXCL10 after allogeneic renal transplantation^21^. In this work, MCMV infection was associated with IL-6, CCL20, CXCL10, IL-21, and CXCL1 after allogeneic transplantation, expanding the list of cytokines and chemokines induced by CMV infection. Observations in clinical transplantation resemble those of the rodent models. Among renal transplant patients, higher plasma levels of chemokines IL-8, MIP-1α and MCP-1 are associated with active CMV infection, and treatment with ganciclovir reduces plasma levels of these chemokines^70^. However, the mechanisms by which MCMV induces production of these inflammatory mediators were not determined in these experiments. Future work may provide additional insights into MCMV-induced dysregulation of the allograft microenvironment during acute rejection.

In this model, MCMV infection promotes the development of Th17 cells with Th1 characteristics. Th1/17 cells have been described in inflammatory diseases including murine experimental autoimmune encephalitis and colitis, and among patients with Candida albicans infection, systemic lupus erythematosus, and Crohn’s disease^71, 72, 73, 74, 75^. Here we report that MCMV-specific Th1/17 cells infiltrate D+R+ allografts and co-express the antiviral cytokine, IFN-γ, but not TNF-α, whereas viral antigen independent, TNF-α and IFN-γ co-expressing Th1/17 cells are also increased by MCMV infection. To our knowledge, this is the first description of MCMV-specific and cytokine-induced Th1/17 cells arising during a pathologic inflammatory disease. It is also notable that both Th17 and Th1 cells are abundant in MCMV-infected allografts. Although conditions promoting Th17 cell development can inhibit Th1 cell differentiation^76, 77, 78, 79, 80^, Th17 cells can also promote accumulation of Th1 cells in inflammatory diseases such as experimental tuberculosis and experimental autoimmune encephalitis, consistent with findings in this model of acute rejection^81, 82^. Transcriptional profiling of MCMV-infected allografts identified the Th1 pathway as one of the most highly upregulated pathways observed in allografts, consistent with TCMR in clinical transplantation^52, 53, 54, 83^. Inhibition of IL-17A reduces neutrophil infiltrates as expected, but unexpectedly also reduces total Th1 cell infiltrates but not virus-specific Th1 cells. This finding suggests the possibility that inhibition of IL-17A may reduce pro-inflammatory Th17 and Th1 cell subsets without impairing intragraft or systemic antiviral Th1 cell responses protecting the host against CMV disease. Further studies are needed to define the interaction between Th17 and Th1 cells in CMV infected allografts, and to characterize the effects of anti-IL-17A upon viral replication and antiviral host immunity. Clinical studies have previously shown that Th17 cells are associated with acute rejection and late allograft dysfunction in populations with high CMV seroprevalence^31, 34^, but CMV was not investigated in these studies. Our work confirms that Th17 cell associated cytokines and transcription factors are elevated during clinical acute rejection, and furthermore shows that AR patients with CMV DNAemia have higher serum quantities of both IL-17A and IFN-γ, but not TNF-α, compared to those without CMV reactivation. This cytokine profile strongly resembles that observed in the MCMV transplant model (Figure 1D) and supports the relevance of findings in the animal model to events occurring during clinical rejection. Further studies among CMV seropositive renal transplant patients are needed to define the specific interactions between CMV reactivation and Th17 cell activity, and their contribution to allograft dysfunction.

There are limitations to this study. For all assays, murine transplants were analyzed at only one timepoint in order to discern early events contributing to development of histopathologic allograft damage at day 14 post-transplant^22^. However, due to the technical complexity of murine microvascular renal transplant surgery, only a limited number of experiments can be performed, so experiments were standardized to one timepoint in order to permit direct comparison of results between experiments. Because of this, the temporal kinetic of MCMV induced chemokine expression, Th17 cell recruitment, and Th1 cell activation were not captured in the current study. We also did not assess developmental plasticity between Th17 and Th1 cell subsets during the course of acute rejection, and it is possible that transdifferentiation between Th17 and Th1 cells could contribute to the Th1/17 cytokine co-expression profiles observed in these studies. In addition, these studies also focused only upon events occurring during acute T cell mediated rejection in CMV R+ recipients. CMV-induced Th17 cell activation was not examined during subclinical CMV DNAemia in the absence of AR, or after primary CMV infection (D+R-transplants). Finally, although the renal transplant population evaluated in this study has nearly universal CMV seropositivity, CMV D/R serostatus was not specifically assessed for these patients. Therefore, it is possible that some patients in this study had CMV D-R- serostatus and consequently had no viral detection. However, this serostatus would be predicted to be distributed similarly among the groups with and without AR, and therefore would be less likely to confound the study results.

In summary, MCMV infection induces recruitment of Th1/17 cells into infected renal allografts, which arise in response to viral antigens as well as via chemokine-induced pathways, and are associated with intragraft IL-17A and IFN-γ. MCMV-induced Th17 cells may contribute to graft injury via recruitment of neutrophils and Th1 cells, with concomitant inhibition of Tregs. Among renal transplant recipients, an imbalance of Th17: Treg cells was observed during acute rejection and Th1/17 cytokines were associated with CMV DNAemia. These findings identified several pathways by which CMV exacerbates allograft inflammation during acute rejection and raises the possibility that inhibition of CMV-induced cytokines might ameliorate allograft injury without adversely impacting protective systemic host antiviral Th1 cell responses. Further studies are needed to investigate such potential interventions against CMV “indirect effects” to advance the goal of mitigating CMV-associated allograft loss in renal transplantation.

## METHODS

### Mice and Virus

BALB/c, C57BL/6, B6.Cg-Tg(TcraTcrb)425Cbn (OT-II) and C57BL/6-*Il17a*^tm1Bcgen^ (*IL17a*-GFP) mice (The Jackson Laboratory, Bar Harbor ME) were housed under specific pathogen-free conditions in animal facilities maintained by the Animal Resources Core (ARC) of The Abigail Wexner Research Institute at Nationwide Children’s Hospital (AWRI) which is accredited by the Assessment and Accreditation of Laboratory Animal Care (AAALAC). Murine experimental protocols were approved by the AWRI Institutional Animal Care and Use Committee. Murine Cytomegalovirus (MCMV) Smith strain with a deletion mutation of the m157 open reading frame (MCMVΔm157) was propagated as described and 1 × 10^6^ plaque forming units (pfu) used to infect donor and recipient mice at least 8 weeks prior to transplantation as described^22^.

### Murine renal transplantation

Allogeneic renal transplantation was performed using male and female BALB/c donors (D) and C57BL/6, *IL-17a*-GFP, or OT-II recipients (R). Syngeneic transplantation was performed between C57BL/6 donors and recipients. Native kidneys were retained, and recipients were treated with cyclosporine (Novartis Pharmaceuticals, East Hanover, NJ) at 10mg/kg/day, subcutaneously once daily until terminal sacrifice.

### Preparation of MCMV Lysate

MCMV lysates were prepared by high speed centrifugation method. In brief, MCMVΔm157 infected, p53 deficient (p53-/-) mouse embryogenic fibroblasts (gift of W. Britt, University of Alabama at Birmingham, Birmingham AL) were lysed by 2 cycles of freeze-thaw and cellular debris pelleted by low speed centrifugation at 2000 x g (Sorvall Legend XTR, Thermo Scientific) for 10 minutes at 4°C. Supernatants were subjected to high speed centrifugation at 16,000 x g for 2 hours at 4°C (Sorvall LYNX 6000, Thermo Scientific). The virus pellet was suspended in 0.9% NaCl and sonicated on ice at 20kHz frequency and 70% amplitude for 15 seconds x 3 cycles (FB505, Fisher Scientific). The protein content of the viral lysate was quantified using BCA protein assay kit (Thermo Scientific, Waltham, MA), aliquoted and stored at -80°C.

### Intracellular cytokine staining (ICS)/Flow cytometry

Splenocytes and intragraft leucocytes were isolated from mice as described previously ^22^. Single leucocyte suspensions were stimulated with either phorbol 12-myristate 13-acetate (PMA)-ionomycin or 5 µg/ml of MCMV peptide pool (Table S3) for 6 hours with brefeldin A (BD Biosciences, San Jose, CA). After stimulation, cells were washed, blocked with anti-CD16/32, and stained for surface markers (Table S5) for 40 minutes protected from light. For chemokine receptor staining, the cells were first stained with anti-mouse CCR6 and CXCR3 at 37°C for 20 minutes prior to stimulation. All other surface stains were performed after stimulation. Fixable viability stain 780 (BD) was added to exclude dead cells. Cells were fixed and permeabilized using True Nuclear Transcription Factor Buffer Set and intracellular cytokine staining (ICS) was performed at 4°C for 1 hour. Isotype controls were included to assess for nonspecific binding of antibodies against intracellular cytokines and transcription factors. For tetramer staining, cells were stained with 3.5 µg/ml of tetramer for 2 hours at 37°C, post stimulation. MHC-I-A^b^ huCLIP staining control was used at identical concentration to assess the nonspecific binding of tetramers. Stained cells were acquired in a LSRII Fortessa (BD) or Attune NxT (Invitrogen, Waltham, MA) and analyzed using FlowJo V10 software (BD).

All antibodies were purchased from Biolegend, except CD45, FoxP3 and IL-10 from BD Bioscience and MHC II (I-A/I-E) from Tonbo Biosciences (San Diego CA). APC labelled I-A^b^-AAHAEINEA329-337 (chicken ovalbumin) and I-A^b^-PVSKMRMATPLLMQA87-101 (human CLIP) tetramers were synthesized by the NIH tetramer Core Facility (Atlanta, GA). The following antibodies were used: CD45 (Clone 30-F11), MHC-II (Clone M5/114.15.2), CD3 (Clone 17A2), CD4 (Clone GK1.5), CD8a (Clone 53-6.7), CXCR3 (Clone CXCR3-173), CCR6 (Clone 29-2L17), CD11b (Clone M1/70), Ly6G (Clone 1A8), IL-17A (Clone TC11-18H10.1), IL-10 (Clone JES5-16E3), IFN-γ (Clone XMG1.2), TNF-α (Clone MP6-XT22) and Foxp3 (Clone MF-23).

### RNA isolation, purification, preparation of library and RNA-Seq analysis

Total RNA from D+ and D- allografts or kidneys of MCMV infected- and uninfected-BALB/c mice was isolated and purified using the RNeasy Mini Kit and RNeasy MinElute Cleanup kit (Qiagen, Germantown MD) according to the manufacturer’s instructions, and purified RNA was treated with DNase (Qiagen) to remove DNA. RNA purity was confirmed using a Nanodrop spectrophotometer (Thermo Fisher, Waltham MA). RNA integrity was assessed using Eukaryote Total RNA Nano Kit on the Bioanalyzer 2100 system (Agilent Technologies, Santa Clara CA). 500 ng of RNA, per sample, was used as input for RNA library preparation. Libraries were generated using the Total RNA TruSeq kit (Illumina, San Diego CA) with Ribo-Zero Complete Globin kit (Illumina) following manufacturer’s recommendations and indices were added to multiplex the data. Final library quality was assessed using the High Sensitivity D1000 kit on the 4200 Tapestation (Agilent). Paired end 150 base pair sequencing was performed on the HiSeq 4000 (Illumina). Each sample was aligned to the GRCm38.p4 assembly of the mouse reference from NCBI using version 2.6.0c of the RNA-Seq aligner STAR ^84^. Transcript features were identified from the GFF file provided with the assembly from NCBI and raw coverage counts were calculated using HTSeq. The raw RNA-Seq gene expression data was normalized and post-alignment statistical analyses were performed using DESeq2 ^85^ and custom analysis scripts written in R. Comparisons of gene expression and associated statistical analysis were made between different conditions of interest using the normalized read counts. All fold change values are expressed as test condition/control condition, where values less than 1 are denoted as the negative of its inverse. Transcripts were considered significantly differentially expressed using a 10% false discovery rate (DESeq2 adjusted p value <= 0.1). The QIAGEN Ingenuity Pathway Analysis software (Qiagen Science, Germantown, MD) was used for canonical pathway analysis and differential gene expression.

### Cytokine and Chemokine detection in tissues by bead immunoassay

Mouse cytokines and chemokines were quantitated in graft and spleen tissue using the LEGENDplex Mouse Inflammation and custom plex panel (Biolegend) according to manufacturer’s instruction ^22^. Spleens and graft tissues were homogenized in SDS free RIPA buffer of pH 8.0 (10mM Tris HCL, 1mM EDTA, 140mM NaCl and 1% Triton X-100) supplemented with protease inhibitor cocktail (Bimake.com, Houston, TX) using a micro-tube homogenizer (Bel-Art, Wayne, NJ). 25 µl of supernatant from each homogenate were processed for the assay according to the manufacturer’s instructions. Samples were acquired in a LSRII Fortessa (BD Biosciences, San Jose, CA) and the data were analyzed using LEGENDplex Data Analysis Software, V8.0 (VigeneTech Inc., Carlisle, MA). Cytokine and chemokine levels were depicted as picograms/gram (pg/g) of tissue.

### In vivo neutralization of IL-17A

Kidney transplant recipients were injected intraperitoneally with 200 µg of neutralizing rat anti-mouse IL-17A or isotype matched control antibodies (BioXCell, West Lebanon, NH) at post-transplant days 1, 3, 5, 7, 9, 12 as indicated through day of terminal sacrifice (day 7 or 14). Allograft kidneys, spleens and peripheral blood were isolated for flow cytometry and histology. Formalin fixed, paraffin embedded allografts were sectioned at 3µm, stained with hematoxylin and eosin, and reviewed by a veterinary renal pathologist (R. C.) blinded to sample identity. A previously published grading scale ^22^ was modified to reflect contemporary rejection criteria as shown in Table S7 ^86, 87^. Scores (0-3) were assigned for 12 histopathologic criteria, for a maximum score of 36.

### Human renal transplant recipients

A retrospective cohort study was conducted using clinical data and samples derived from a previously completed prospective observational cohort study that was approved by the Institutional Review Board (IRB) and conducted at the Department of Renal Transplant Surgery of the Post Graduate Institute of Medical Education and Research (PGIMER) in India to investigate the genetic, phenotypic and functional variations of T and B cells at the time of renal allograft rejection ^88, 89, 90^. For the parent study, adults (age > 18 years) diagnosed with end stage renal disease undergoing first renal transplantation between July 2012 and August 2013 were included in the study after obtaining their written informed consent. Exclusion criteria were recipients of multiorgan or prior organ transplant and known HIV, hepatitis B or C infection. The recipients were followed for 12 months post-transplant. Demographic and clinical data were collected: donor and recipient age and sex, recipient’s disease history, HLA types, cross match, induction immunosuppression, graft ischemia time, post-transplant creatinine and immunosuppressant levels. For rejection episodes, histopathologic rejection classification and DSA status were collected from the clinical record. Blood was collected from all subjects at pre-transplant, 1-, 3-, 6- and 12-months post-transplant and at the time of rejection. The retrospective cohort study was performed to determine the association between blood mRNA RORγt: FOXP3 ratio and acute renal allograft rejection. The study was approved as exempt research under the category of secondary research using previously collected specimen by the PGIMER institutional review board (Endorsement No. 8766-PG-10-1TRG/8448). Cases of acute allograft rejection (AR) were identified by searching the database. Eligible cases included all the recipients with acute cell mediated allograft rejection (N= 18) and acute mixed allograft rejection (N=6). A control cohort of 29 recipients without acute rejection were matched with the cases for age and sex.

### RNA extraction, cDNA preparation and Gene expression analysis in human transplant recipients

Total RNA was extracted using QIAamp RNA Blood Mini Kit (Qiagen, Germantown, MD). First strand cDNA was synthesized from 200 ng of total RNA and a mixture of Oligo (dT) and random hexamer primers in a 20 μl reaction volume using RevertAid first strand cDNA synthesis Kit (Thermofisher Scientific, Waltham, MA). PSurity of the cDNA was measured by A260/A280 ratio using UV spectroscopy (Thermofisher Scientific, Waltham, MA). Relative gene expression of RORγt and FOXP3 was performed using Real Time PCR (Lightcycler 480, Roche Molecular System, Pleasanton, CA), SYBR Green and primers specific to RORγt, FOXP3 and β-actin (Table S6) ^91^. The amplification protocol consisted of 95°C for 10 minutes, followed by 45 cycles of 95°C for 10 seconds, 60°C for 15 seconds and 72°C for 20 seconds. Relative gene expression level (Fold Change) was calculated using the 2^-ΔΔC^_T_ method and the quantitation was normalized to endogenous gene expression of β-actin.

### Detection of cytokines in human peripheral blood

Peripheral blood in serum blood collection tubes were allowed to clot at room temperature for 30 minutes, centrifuged at 1500rpm for 10 minutes to collect the serum and store in aliquots at -80°C until further use. Cytokines in the serum sample were measured using human Th1/Th2/Th17 cytometric bead array kit (BD Bioscience, San Jose, CA) according to the manufacture’s instruction. Briefly, 50µl of serum or serially diluted standard cytokines cocktail were mixed with 50µl of capture beads and 50µl of Th1/Th2/Th17 PE detection reagent. The mixtures were incubated for 3 hours at room temperature at dark, washed and acquired in BD FACS Aria II (BD Biosciences, San Jose, CA) with 488nm and 633nm lasers using BD FACSDiva software version 6.0. Cytokine level in the peripheral blood were expressed as picograms/ milliliter (pg/ml) of serum.

### HCMV viral load in peripheral blood by quantitative real time PCR

Peripheral blood was collected in BD Vacutainer EDTA tubes. Genomic DNA was extracted from the blood using QIAamp DNA blood mini kit (Qiagen Science, Germantown, MD). The CMV viral load was performed as described previously^92^. Briefly, HCMV glycoprotein B gene (gB) in the DNA sample was amplified in Lightcycler 480 real time PCR platform (Roche Molecular System, Pleasanton, CA) using TaqMan probe (LightCycler TaqMan Master, Roche Molecular System, Pleasanton, CA) and primers specific to HCMV gB (Table S6) ^92^. The amplification protocol consisted of 95°C for 10 minutes, followed by 45 cycles of 58°C for 10 seconds and 72°C for 20 seconds. Viral copies/ml of peripheral blood was calculated using the standard curve generated with serially diluted cloned DNA standard plasmids (Roche Molecular System, Pleasanton, CA).

### Statistics

Descriptive analyses were used to summarize patients’ demographic characteristics using ranges and frequency distributions as appropriate. Results are presented as mean ± standard deviation (SD) unless otherwise indicated. Continuous variables were analyzed using the two-tailed Student’s t test or the one-way analysis of variance. Categorical variables were compared using the Chi-square test. Statistical analyses were performed using GraphPad Prism 8 (San Diego, CA) accepting a p value less than 0.05 statistically significant.

## DATA AVAILABILITY

RNA sequencing data have been deposited in NCBI’s Gene Expression Omnibus^93^ and are accessible through GEO series accession number GSE179788.

## ACKNOWLEDGEMENTS

We are grateful to the renal transplant patients for their participation in this study. We thank Sonya Maher, Jessica Ghere, Irina Kaptsan, Brianna Graber, and Kaitlyn Flint for providing technical assistance in murine experiments. Funding for this work was provided by NIH R01AI101138 (M.S.), the Abigail Wexner Research Institute at Nationwide Children’s Hospital (M.S.) and the intramural Research grant of the Post graduate institute of Medical Education and Research (M.M. and R.W.M)

## AUTHOR CONTRIBUTIONS

M.S conceptualized the study. M.S., R.D., V.M.V., M.M. and R.W.M., designed the experiments. R.D., S.A., and S.R.B. performed the experiments. M.M., A.S., and Q.Z. performed renal transplant surgery. R.D., M.S., R.C., and J.R.F., analyzed the data. R.D., and M.S., wrote the manuscript. All authors edited the manuscript.

## SUPPLEMENTAL INFORMATION TITLES AND LEGENDS

**Figure S1.**
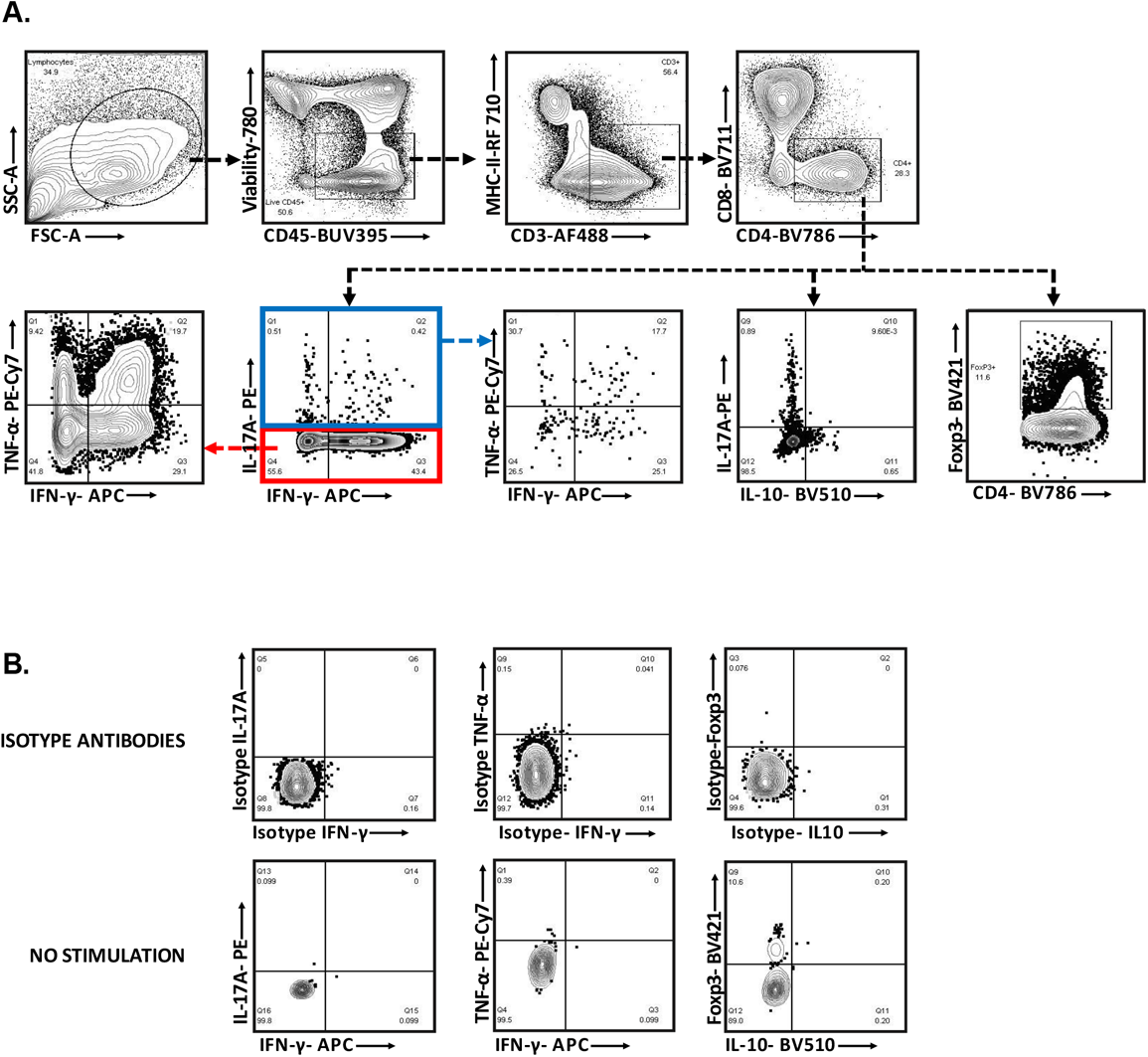
Flow cytometry gating strategy. (A) Gating strategy for cytokine expressing Th1, Th17, and Treg cells. Lymphocytes were first identified by a forward scatter (FSC) and side scatter (SSC) gate. Viable CD45^+^ cells were gated and CD4^+^ T cells were identified within CD3^+^MHCII^-^ gate. Th17 cells were identified as IL-17A expressing CD4^+^ T cells. IL-17A^-^IFN-γ^+^/TNF-α^+^ CD4^+^ T- cells were identified as Th1 cells. FoxP3^+^ CD4^+^ T- cells were identified as Tregs. (B) Isotype staining to detect the nonspecific binding of antibodies against intracellular cytokines and Foxp3. The no-stimulation condition was included in the assay to gate the endogenous fluorescence.

**Figure S2.**
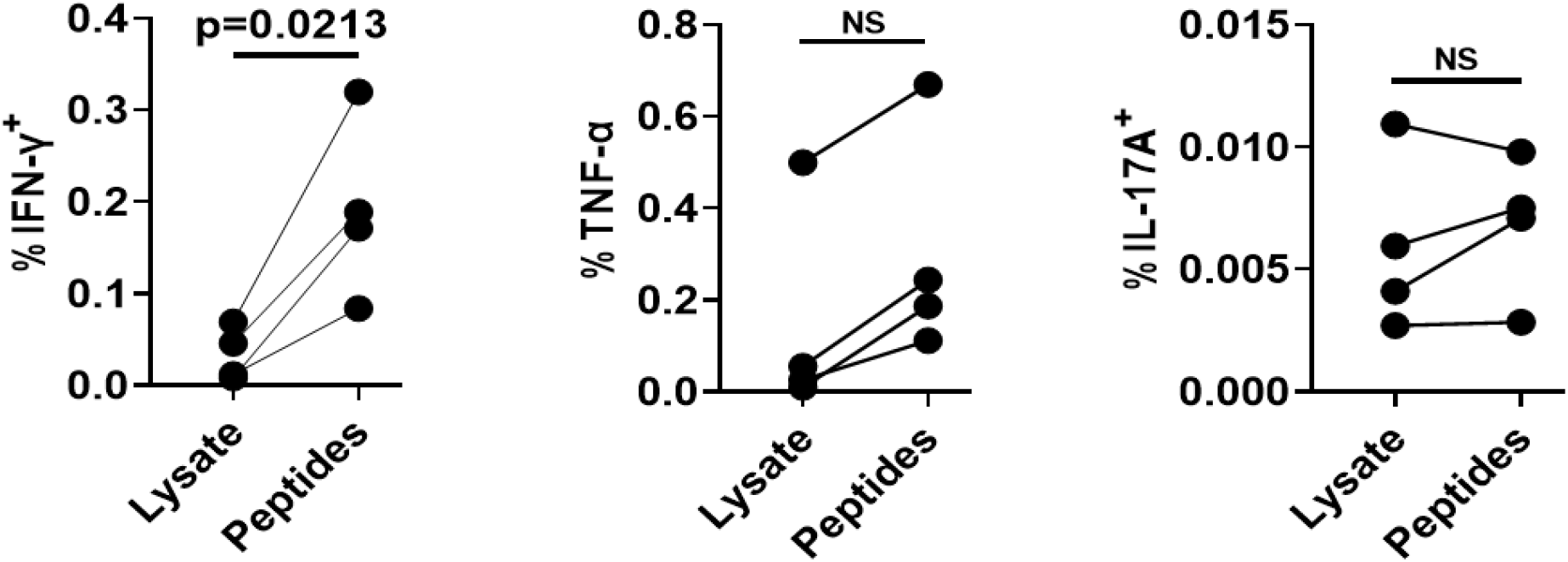
Comparison of MCMV-specific CD4^+^ T cell responses after stimulation with viral lysate or MCMV peptides. Spleenocytes from D+R+ transplants were stimulated with MCMV lysate or a pool of 15 class II-restricted MCMV peptides (Table S3) and then stained for IFN-γ, TNF-α and IL-17A expressing CD4^+^ T cells. Frequencies of CD4^+^ T cells that expressed cytokines when stimulated with lysate or peptides. The differences between the two groups were analyzed using two- sided Student’s t test. NS, not significant (p>0.05).

**Figure S3.**
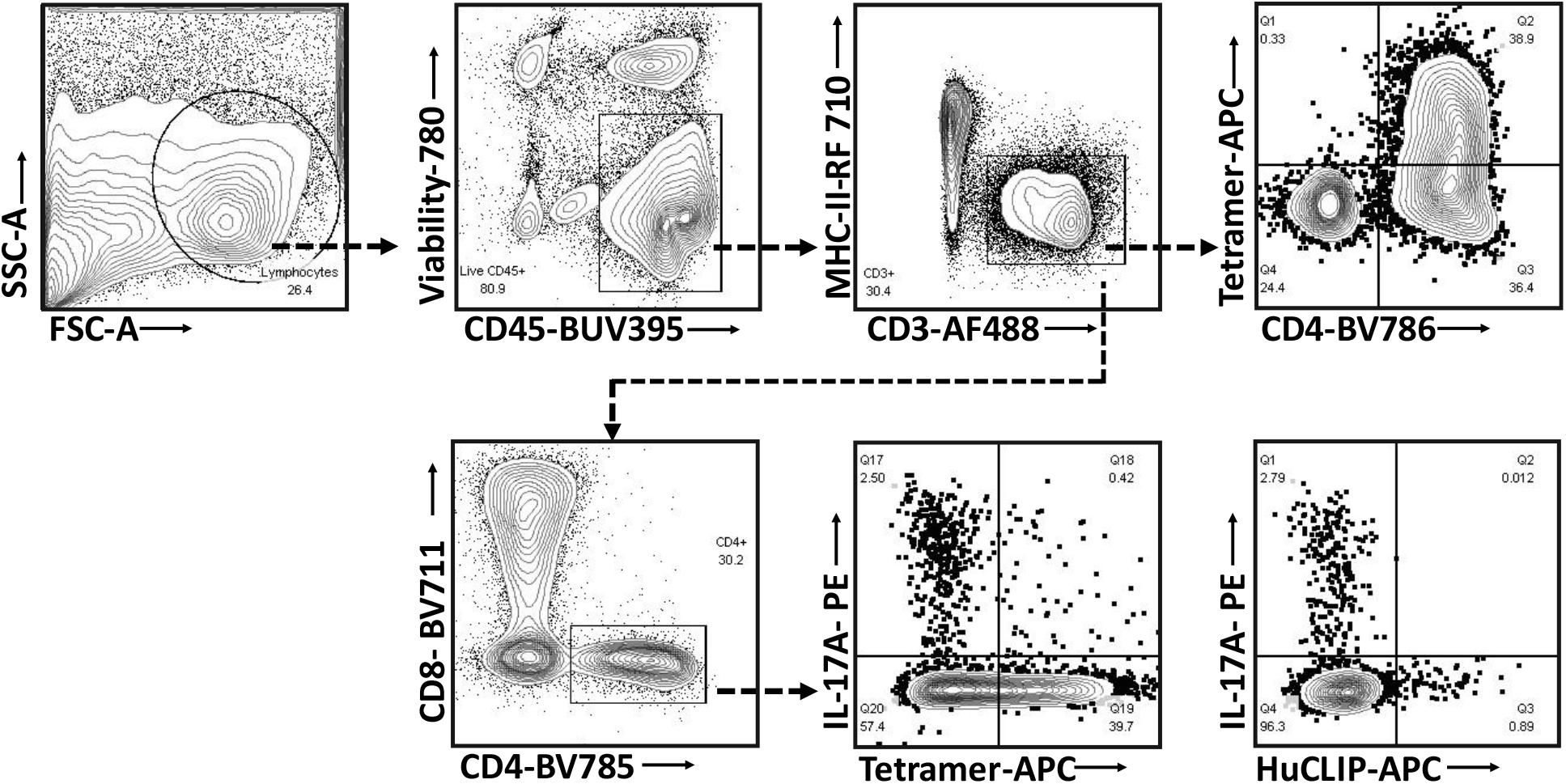
Flow cytometry gating strategy for I-A^b^-OVA_323-339_-APC tetramer^+^ cells. CD3^+^MHCII^-^ T- cells were gated as described previously. I-A^b^-OVA_323-339_-APC tetramer^+^ CD4^+^ T- cells were identified within CD3^+^MHCII^-^ T- cells and IL-17A^+^ Tetramer^+^/^-^ cells were gated within CD4^+^ T- cells. HuCLIP-APC tetramers were used as control staining for the OVA_323-339_-APC tetramer. HuCLIP, Human MHC Class II-associated invariant chain.

**Figure S4.**
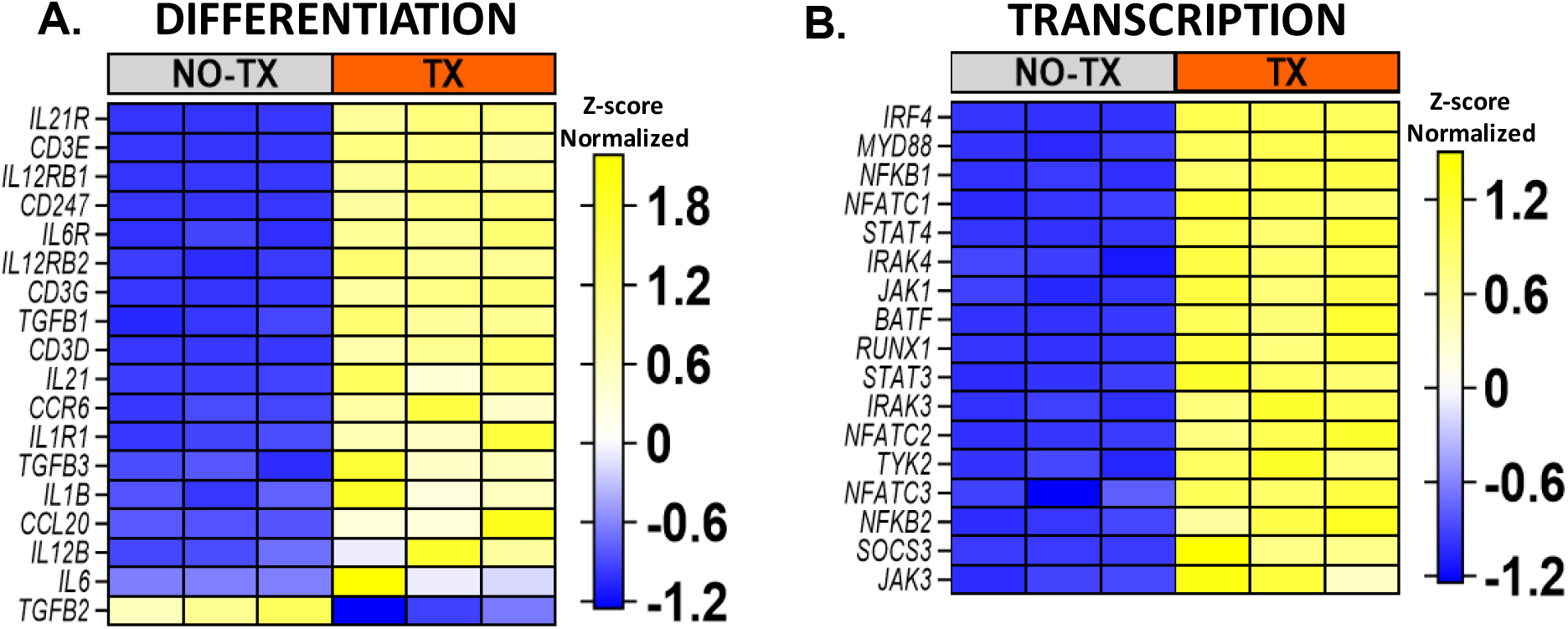
Differential expression of transcripts between D- allografts (TX) and native non-transplant MCMV uninfected kidneys (NO-TX). (A) Differential expression of transcripts involved in Th17 differentiation. (B) Differential expression of transcripts for Th17 cell associated transcription factors.

**Figure S5.**
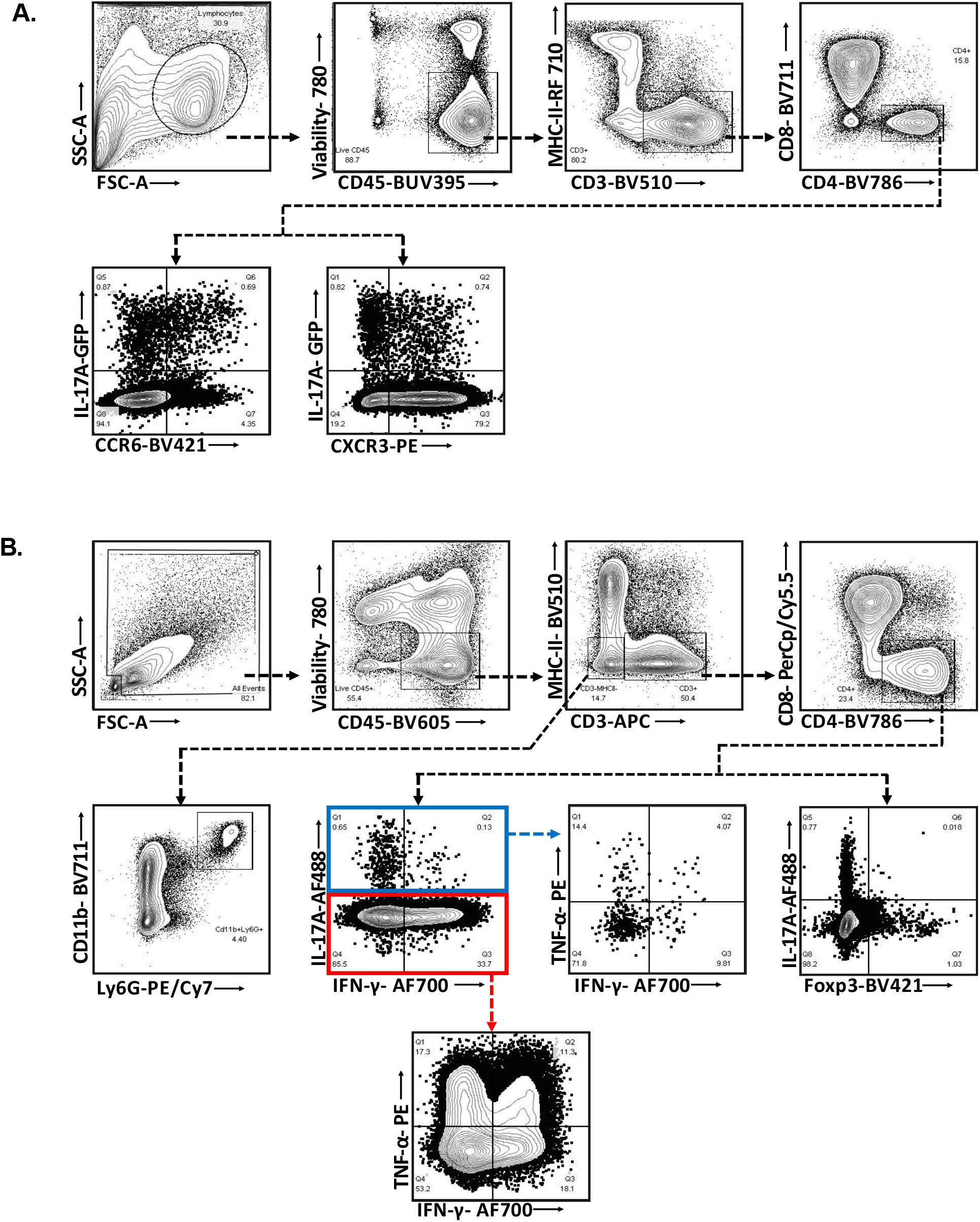
Flow cytometry gating strategy. (A) Expression of chemokine receptors by Th17 cells. CD4^+^ T- cells were gated on live CD45^+^MHCII^-^CD3^+^ T- cells as described previously. CCR6^+^/CXCR3^+^IL-17A^+^ cells were identified within CD4^+^ T- cells gate. (B) Neutrophil and cytokine expressing CD4^+^ T- cells in anti-IL-17A treated and control transplant mice. CD11b^+^ Ly6G^+^ Neutrophils were gated on live CD45^+^MHCII^-^CD3^-^ populations. Th1, Th17 and Tregs were gated as described previously.

**Figure S6.**
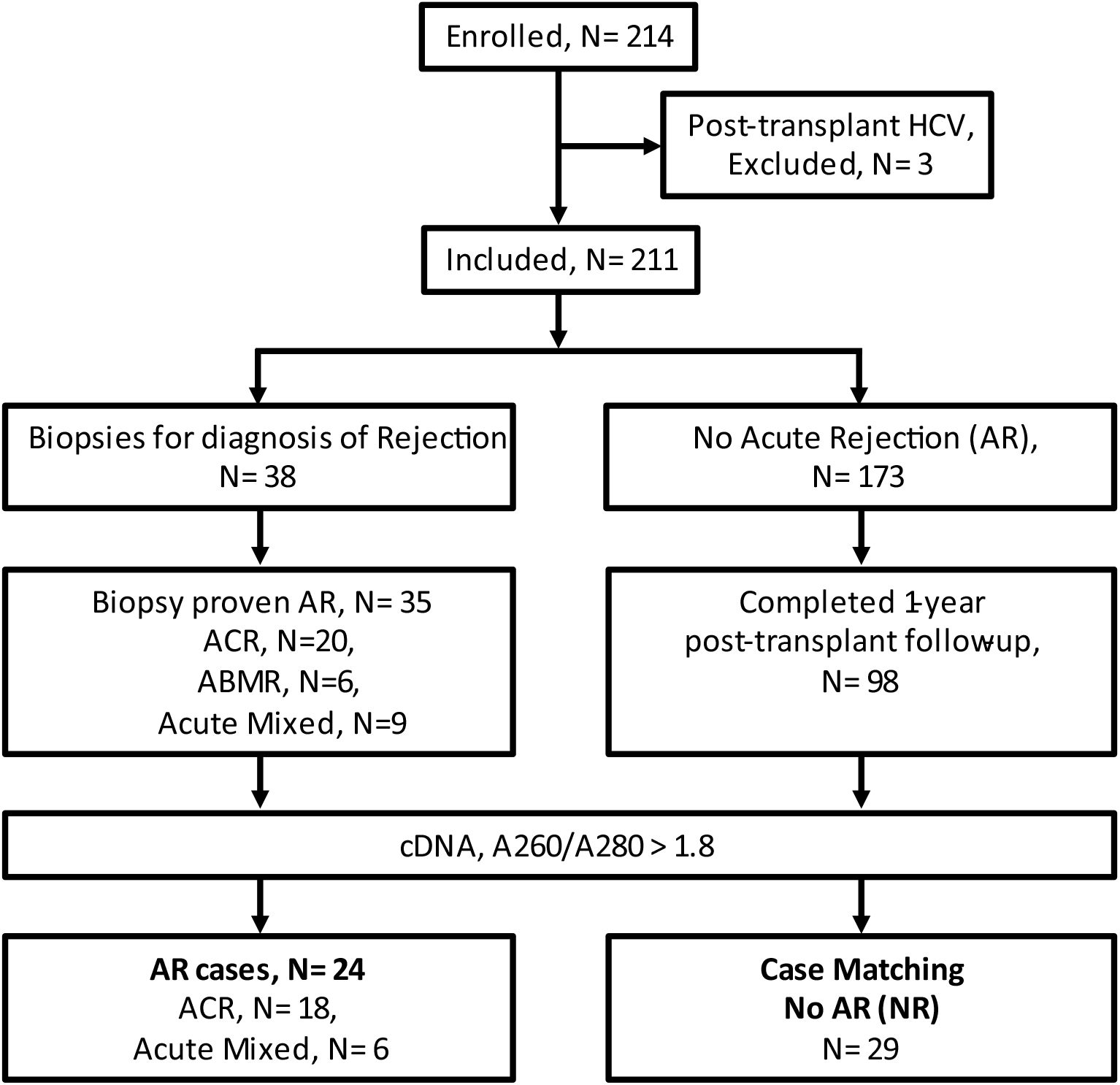
Human Study population. Of 214 enrolled patients, 3 were excluded due to post-transplant HCV infection. Of the 211 remaining patients, 35 had biopsy proven acute rejection, of whom 24 patients had blood RNA samples of sufficient quality for RT-PCR analysis (AR group). Of the 173 patients without AR, 29 patients (NR group) with blood RNA of sufficient quality for RT-PCR were matched by age and sex to patients in the AR group. HCV, Hepatitis- C virus; ACR, Acute Cellular Rejection; ABMR, Antibody mediated rejection; cDNA, complimentary DNA; AR, Acute Rejection.

**Table S1.** Demographic details of Renal Transplant Recipients.

| Variables | No rejection (NR),<br>N= 29 | Acute Rejection (AR),<br>N=24 |
| --- | --- | --- |
| Mean Age in Years (Range) | 38.38 ± 12.13 (20- 58) | 32.33 ± 7.91 (12-44) |
| Gender |  |  |
| Male | 24 (82.75%) | 20 (83.34%) |
| Female | 5 (17.24%) | 4 (16.66%) |
| Underlying disease for ESRD |  |  |
| CGN-CRF | 14 (48.27%) | 9 (37.50 %) |
| Hypertension | 7 (24.13%) | 9 (37.50 %) |
| Diabetic Nephropathy | 3 (10.34%) | 1 (4.16%) |
| IgA Nephropathy | 1 (3.44%) | 2 (8.33%) |
| Others<sup>a</sup> | 3 (10.34%) | 3 (12.50%) |
| Allograft |  |  |
| Living related | 13 (44.80%) | 14 (58.33%) |
| Living Unrelated | 11 (37.93%) | 6 (25.00%) |
| Deceased | 5 (17.24%) | 4 (16.66%) |
| HLA Mismatch | 4.0 ± 1.9 (0- 6) | 4.0 ± 1.4 (2- 6) |
| Induction Immunosuppression |  |  |
| ATG | 1 (3.44%) | 1 (4.16 %) |
| Anti IL2R | 4 (13.79%) | 5 (20.83%) |
| None | 24 (82.75%) | 18 (75.00%) |
| Maintenance Immunosuppression |  |  |
| Tac/MMF/Steroid | 27 (93.10%) | 24 (100%) |
| CsA/MMF/ Steroid | 1 (3.44%) | 0 |
| Tac/AZA/Steroid | 1 (3.44%) | 0 |
| Types of Rejection |  |  |
| ACR | NA | 18 (75.00%) |
| Mixed | NA | 6 (25.00%) |
| Mean onset of Rejection, days | NA | 32.44 ± 49.46 |

Others<sup>a</sup> Alport Syndrome, Autosomal Dominant Polycystic Kidney Disease, Collapsing thrombotic microangiopathy and Obstructive Uropathy. ESRD, End-stage renal disease; CGN-CRF, Chronic glomerulonephritis-Chronic renal failure; HLA, Human leukocyte antigen; ATG, Anti-thymoglobulin; Tac, Tacrolimus; MMF, Mycophenolate Mofetil; CsA, Cyclosporin A; AZA, Azathioprine; ACR, Acute cellular rejection.

**Table S2.** Cases of CMV reactivation in Renal Transplant Recipients.

|  | NR | AR | P value* |
| --- | --- | --- | --- |
| CMV- | 16 | 12 | 0.0159 |
| CMV+ | 1 | 8 |  |
| Total | 17 | 20 |  |

CMV, Cytomegalovirus; NR, No Rejection; AR, Acute Rejection. \*Chi-square test.

**Table S3.**
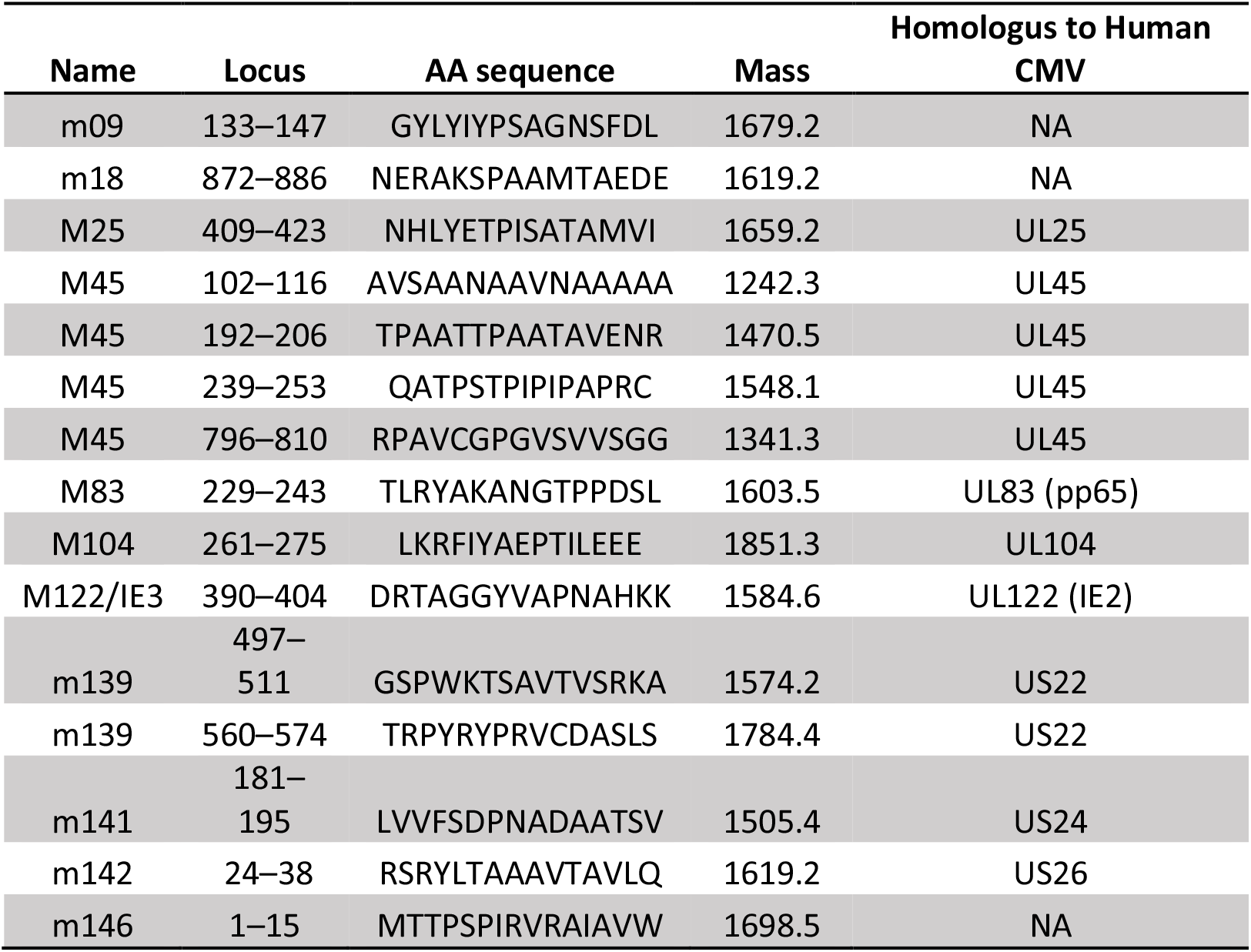
C57BL/6- CD4 specific MCMV Peptides.

**Table S4.**
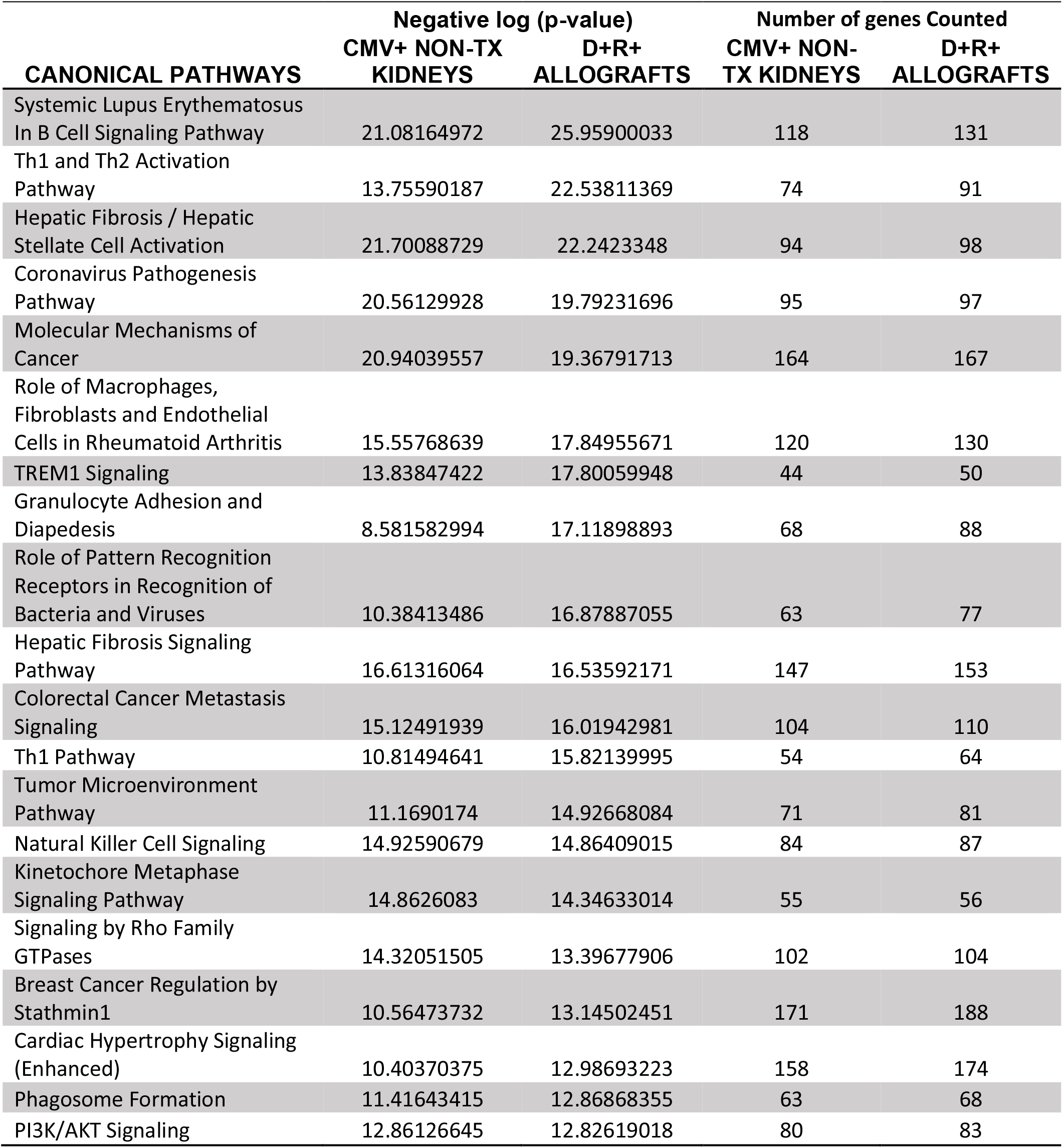
Top 20 Canonical Pathways.

**Table S5.**
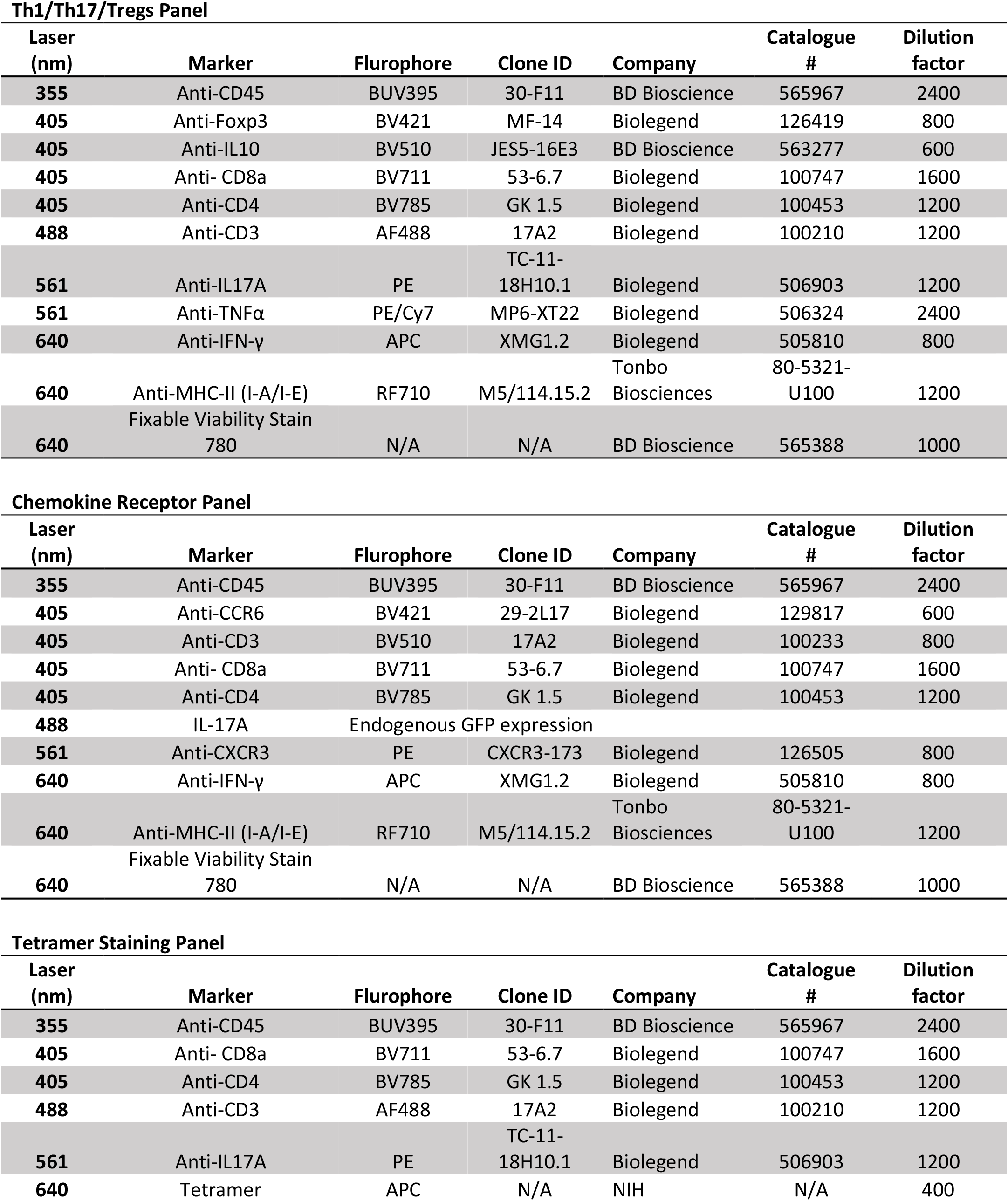
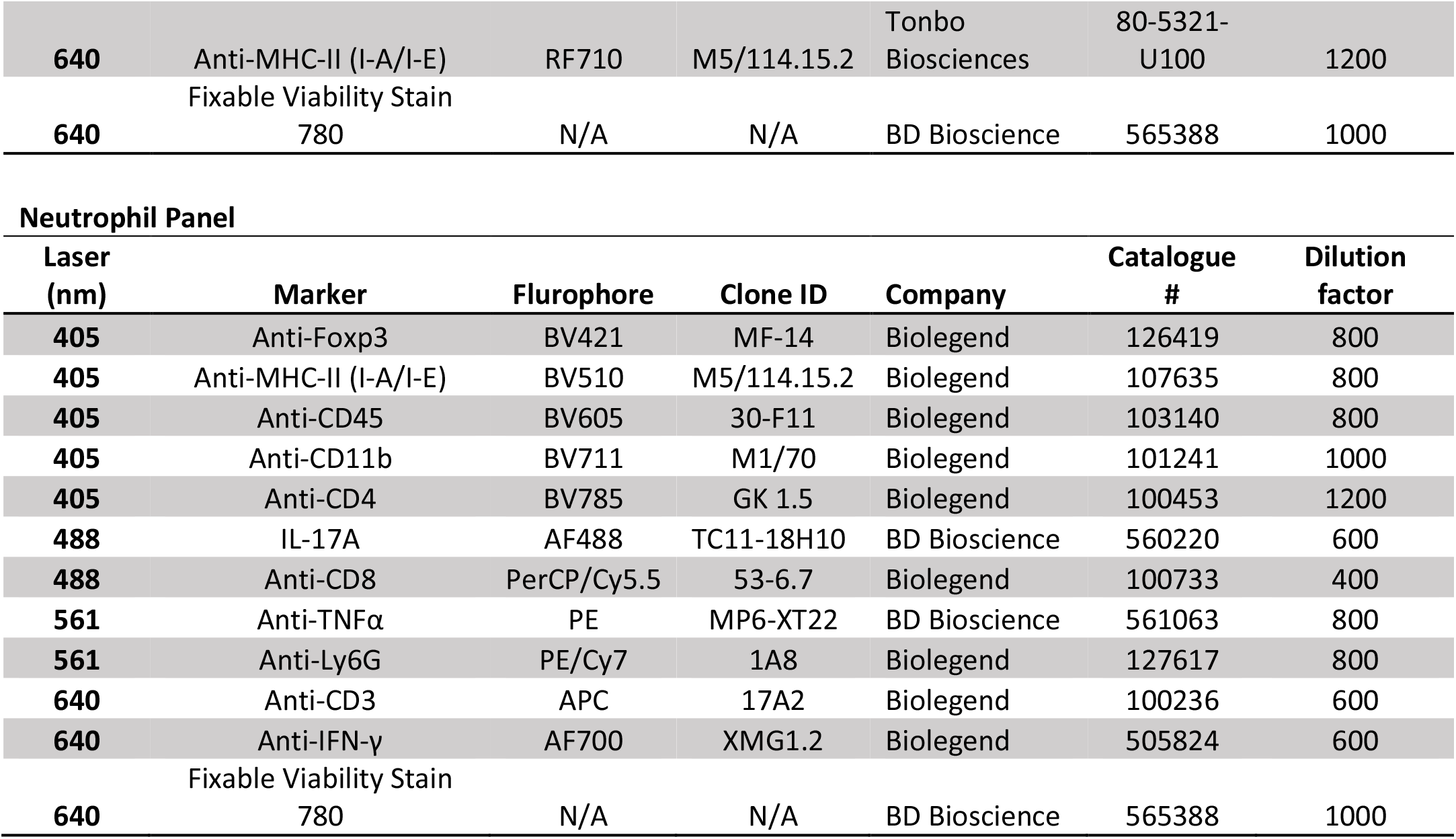
Flow cytometry panels.

**Table S6.**
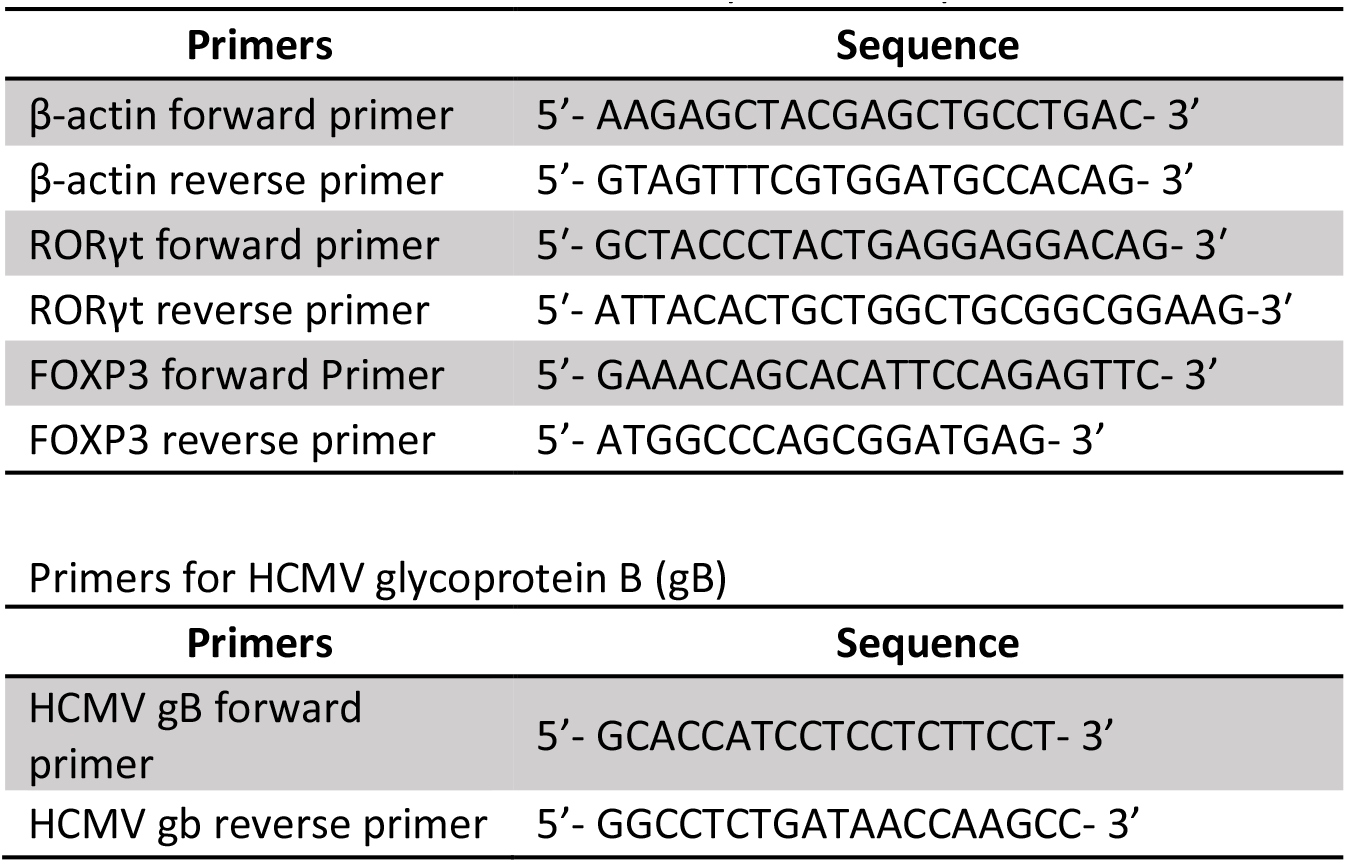
Primers for FOXP3 and RORγt mRNA expression.

**Table S7.**
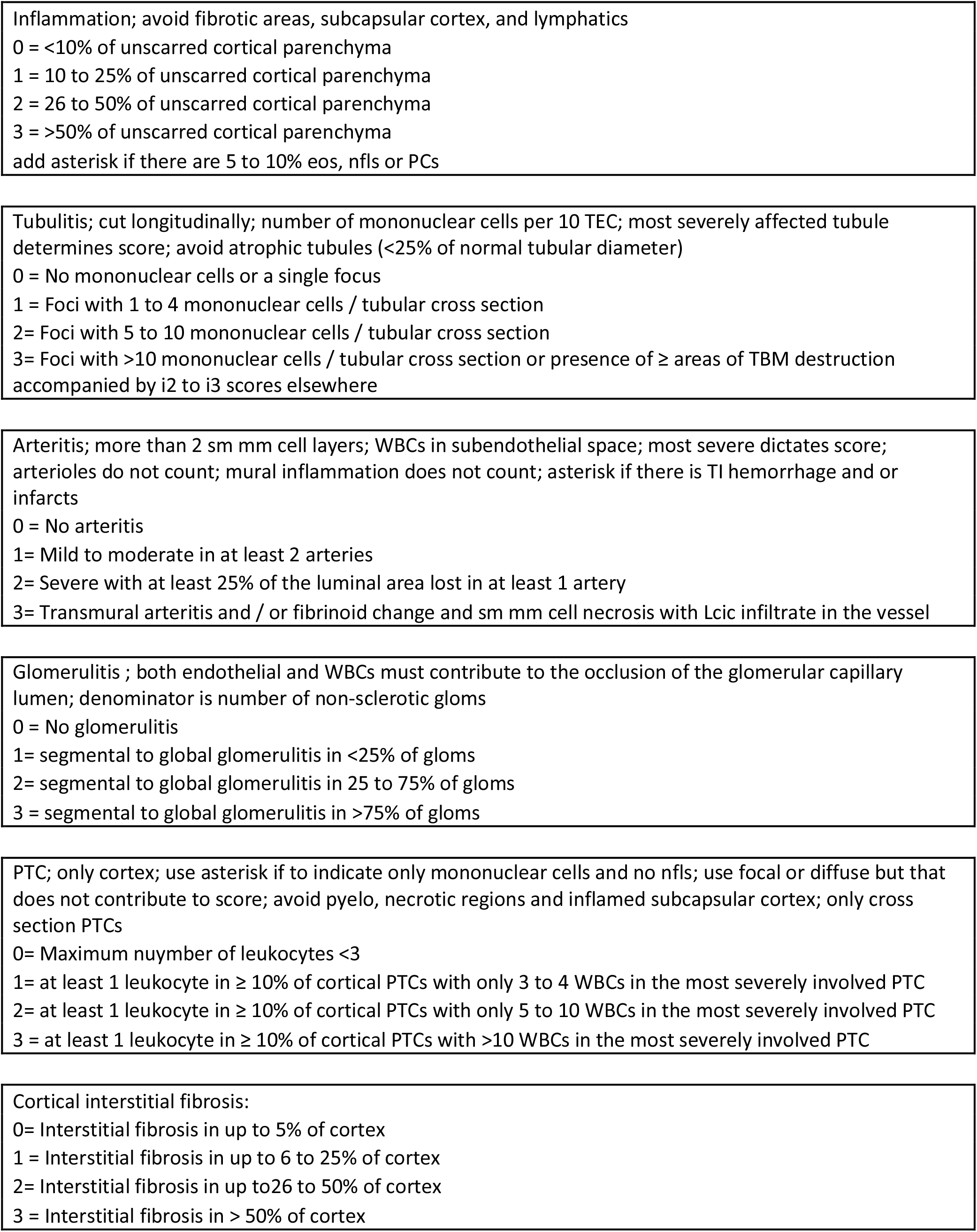
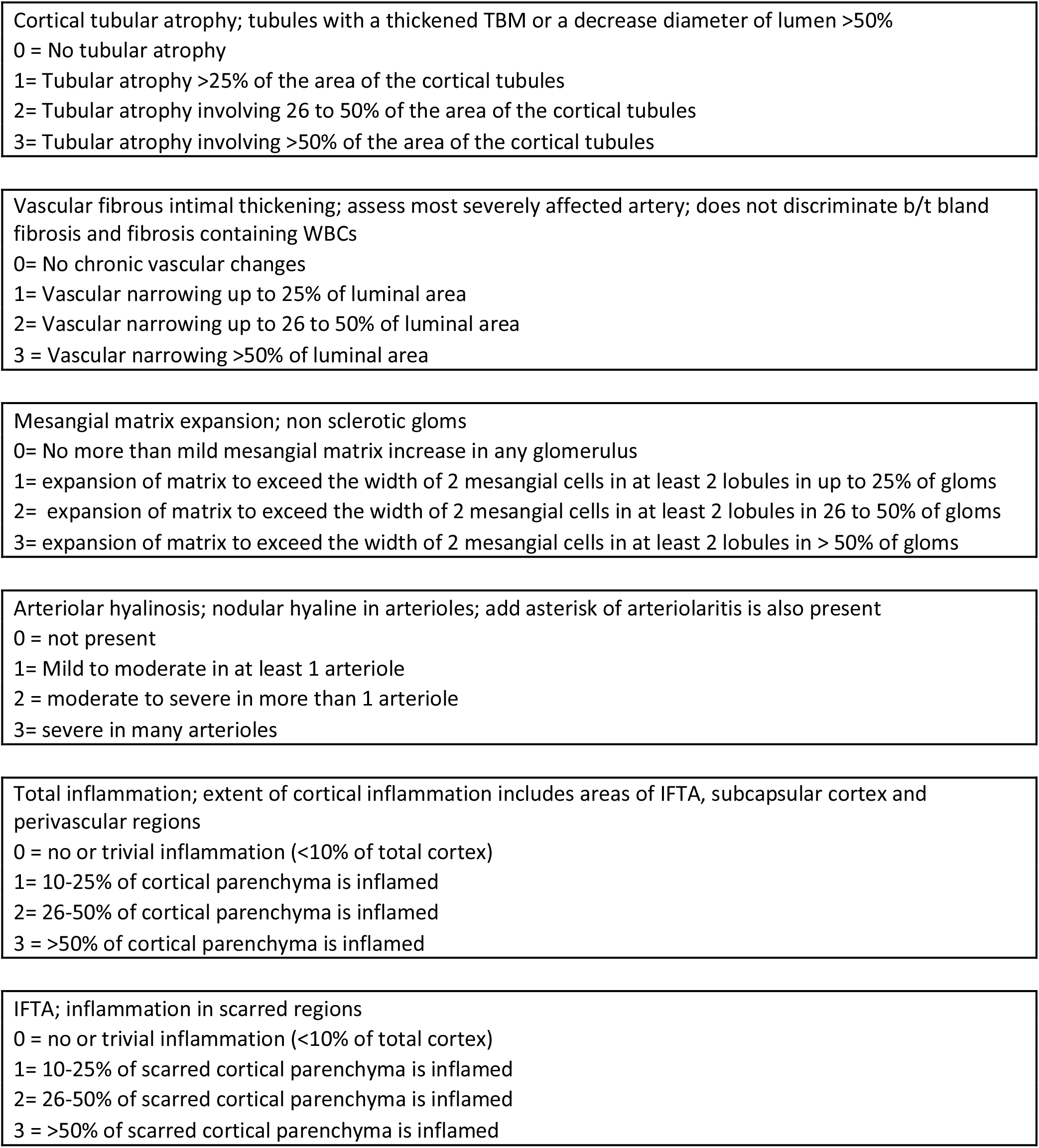
Histology Scoring Criteria.

## REFERENCES

1. Das, B., Kaur, G. & Basu, S. Seroprevalence of cytomegalovirus antibodies among blood donors and Multitransfused recipients--a study from north India. Transfus Apher Sci 50, 438–442 (2014).

2. Lachmann, R. et al.. Cytomegalovirus (CMV) seroprevalence in the adult population of Germany. PLoS One 13, e0200267 (2018).

3. Warnecke, J.M. et al.. Seroprevalences of antibodies against ToRCH infectious pathogens in women of childbearing age residing in Brazil, Mexico, Germany, Poland, Turkey and China. Epidemiol Infect 148, e271 (2020).

4. Myerson, D., Hackman, R.C., Nelson, J.A., Ward, D.C. & McDougall, J.K. Widespread presence of histologically occult cytomegalovirus. Hum Pathol 15, 430–439 (1984).

5. Hendrix, R.M., Wagenaar, M., Slobbe, R.L. & Bruggeman, C.A. Widespread presence of cytomegalovirus DNA in tissues of healthy trauma victims. J Clin Pathol 50, 59–63 (1997).

6. Klenerman, P. & Oxenius, A. T cell responses to cytomegalovirus. Nat Rev Immunol 16, 367–377 (2016).

7. Freeman, R.B., Jr. The ‘indirect’ effects of cytomegalovirus infection. Am J Transplant 9, 2453–2458 (2009).

8. Humar, A. et al.. Association between cytomegalovirus disease and chronic rejection in kidney transplant recipients. Transplantation 68, 1879–1883 (1999).

9. Tong, C.Y., Bakran, A., Peiris, J.S., Muir, P. & Herrington, C.S. The association of viral infection and chronic allograft nephropathy with graft dysfunction after renal transplantation. Transplantation 74, 576–578 (2002).

10. Toupance, O. et al.. Cytomegalovirus-related disease and risk of acute rejection in renal transplant recipients: a cohort study with case-control analyses. Transpl Int 13, 413–419 (2000).

11. Gamadia, L.E., Rentenaar, R.J., van Lier, R.A. & ten Berge, I.J. Properties of CD4(+) T cells in human cytomegalovirus infection. Hum Immunol 65, 486–492 (2004).

12. Jeitziner, S.M., Walton, S.M., Torti, N. & Oxenius, A. Adoptive transfer of cytomegalovirus-specific effector CD4+ T cells provides antiviral protection from murine CMV infection. Eur J Immunol 43, 2886–2895 (2013).

13. Rentenaar, R.J. et al.. Development of virus-specific CD4(+) T cells during primary cytomegalovirus infection. J Clin Invest 105, 541–548 (2000).

14. Sester, M. et al.. Levels of virus-specific CD4 T cells correlate with cytomegalovirus control and predict virus-induced disease after renal transplantation. Transplantation 71, 1287–1294 (2001).

15. Walter, E.A. et al.. Reconstitution of cellular immunity against cytomegalovirus in recipients of allogeneic bone marrow by transfer of T-cell clones from the donor. N Engl J Med 333, 1038–1044 (1995).

16. Reddehase, M.J., Mutter, W., Münch, K., Bühring, H.J. & Koszinowski, U.H. CD8-positive T lymphocytes specific for murine cytomegalovirus immediate-early antigens mediate protective immunity. J Virol 61, 3102–3108 (1987).

17. Dunn, H.S. et al.. Dynamics of CD4 and CD8 T cell responses to cytomegalovirus in healthy human donors. J Infect Dis 186, 15–22 (2002).

18. Razonable, R.R. & Humar, A. Cytomegalovirus in solid organ transplant recipients-Guidelines of the American Society of Transplantation Infectious Diseases Community of Practice. Clin Transplant 33, e13512 (2019).

19. Witzke, O. et al.. Valganciclovir Prophylaxis Versus Preemptive Therapy in Cytomegalovirus-Positive Renal Allograft Recipients: Long-term Results After 7 Years of a Randomized Clinical Trial. Transplantation 102, 876–882 (2018).

20. Lautenschlager, I. et al.. CMV increases inflammation and accelerates chronic rejection in rat kidney allografts. Transplantation Proceedings 29, 802–803 (1997).

21. Soule, J.L., Streblow, D.N., Andoh, T.F., Kreklywich, C.N. & Orloff, S.L. Cytomegalovirus accelerates chronic allograft nephropathy in a rat renal transplant model with associated provocative chemokine profiles. Transplant Proc 38, 3214–3220 (2006).

22. Li, M. et al.. NK cell and Th17 responses are differentially induced in murine cytomegalovirus infected renal allografts and vary according to recipient virus dose and strain. Am J Transplant 18, 2647–2662 (2018).

23. Saunders, U. et al.. Murine Cytomegalovirus-Induced Complement-fixing Antibodies Deposit in Murine Renal Allografts During Acute Rejection. Transplantation (2020).

24. Shimamura, M. et al.. Ganciclovir prophylaxis improves late murine cytomegalovirus-induced renal allograft damage. Transplantation 95, 48–53 (2013).

25. Korn, T. et al.. IL-21 initiates an alternative pathway to induce proinflammatory T(H)17 cells. Nature 448, 484–487 (2007).

26. Liang, S.C. et al.. Interleukin (IL)-22 and IL-17 are coexpressed by Th17 cells and cooperatively enhance expression of antimicrobial peptides. J Exp Med 203, 2271–2279 (2006).

27. Acosta-Rodriguez, E.V., Napolitani, G., Lanzavecchia, A. & Sallusto, F. Interleukins 1beta and 6 but not transforming growth factor-beta are essential for the differentiation of interleukin 17- producing human T helper cells. Nat Immunol 8, 942–949 (2007).

28. Miossec, P. & Kolls, J.K. Targeting IL-17 and TH17 cells in chronic inflammation. Nat Rev Drug Discov 11, 763–776 (2012).

29. Zambrano-Zaragoza, J.F., Romo-Martinez, E.J., Duran-Avelar Mde, J., Garcia-Magallanes, N. & Vibanco-Perez, N. Th17 cells in autoimmune and infectious diseases. Int J Inflam 2014, 651503 (2014).

30. Crispim, J.C. et al.. Interleukin-17 and kidney allograft outcome. Transplant Proc 41, 1562–1564 (2009).

31. Chung, B.H. et al.. Increase of Th17 Cell Phenotype in Kidney Transplant Recipients with Chronic Allograft Dysfunction. PLoS One 10, e0145258 (2015).

32. Fabrega, E., Lopez-Hoyos, M., San Segundo, D., Casafont, F. & Pons-Romero, F. Changes in the serum levels of interleukin-17/interleukin-23 during acute rejection in liver transplantation. Liver Transpl 15, 629–633 (2009).

33. Fan, H. et al.. Increase of peripheral Th17 lymphocytes during acute cellular rejection in liver transplant recipients. Hepatobiliary & Pancreatic Diseases International 11, 606–611 (2012).

34. Chung, B.H. et al.. Higher infiltration by Th17 cells compared with regulatory T cells is associated with severe acute T-cell-mediated graft rejection. Exp Mol Med 43, 630–637 (2011).

35. Yapici, U. et al.. Interleukin-17 positive cells accumulate in renal allografts during acute rejection and are independent predictors of worse graft outcome. Transpl Int 24, 1008–1017 (2011).

36. Strehlau, J. et al.. Quantitative detection of immune activation transcripts as a diagnostic tool in kidney transplantation. Proc Natl Acad Sci U S A 94, 695–700 (1997).

37. Van Kooten, C. et al.. Interleukin-17 activates human renal epithelial cells in vitro and is expressed during renal allograft rejection. J Am Soc Nephrol 9, 1526–1534 (1998).

38. Woltman, A.M. et al.. MIP-3alpha/CCL20 in renal transplantation and its possible involvement as dendritic cell chemoattractant in allograft rejection. Am J Transplant 5, 2114–2125 (2005).

39. Gorbacheva, V., Fan, R., Li, X. & Valujskikh, A. Interleukin-17 promotes early allograft inflammation. Am J Pathol 177, 1265–1273 (2010).

40. Yuan, X. et al.. A novel role of CD4 Th17 cells in mediating cardiac allograft rejection and vasculopathy. J Exp Med 205, 3133–3144 (2008).

41. Khader, S.A., Gaffen, S.L. & Kolls, J.K. Th17 cells at the crossroads of innate and adaptive immunity against infectious diseases at the mucosa. Mucosal Immunol 2, 403–411 (2009).

42. Yang, B.H. et al.. Foxp3(+) T cells expressing RORgammat represent a stable regulatory T-cell effector lineage with enhanced suppressive capacity during intestinal inflammation. Mucosal Immunol 9, 444–457 (2016).

43. Khader, S.A. & Gopal, R. IL-17 in protective immunity to intracellular pathogens. Virulence 1, 423–427 (2010).

44. Aghbash, P.S. et al.. The role of Th17 cells in viral infections. Int Immunopharmacol 91, 107331 (2021).

45. Lukacs, N.W. et al.. Respiratory virus-induced TLR7 activation controls IL-17-associated increased mucus via IL-23 regulation. J Immunol 185, 2231–2239 (2010).

46. Paquissi, F.C. Immunity and Fibrogenesis: The Role of Th17/IL-17 Axis in HBV and HCV-induced Chronic Hepatitis and Progression to Cirrhosis. Front Immunol 8, 1195 (2017).

47. Arens, R. et al.. Cutting edge: murine cytomegalovirus induces a polyfunctional CD4 T cell response. J Immunol 180, 6472–6476 (2008).

48. Takada, M., Nadeau, K.C., Shaw, G.D., Marquette, K.A. & Tilney, N.L. The cytokine-adhesion molecule cascade in ischemia/reperfusion injury of the rat kidney. Inhibition by a soluble P-selectin ligand. J Clin Invest 99, 2682–2690 (1997).

49. Zhang, Z. et al.. A clinically relevant murine model unmasks a “two-hit” mechanism for reactivation and dissemination of cytomegalovirus after kidney transplant. Am J Transplant 19, 2421–2433 (2019).

50. Harrington, L.E. et al.. Interleukin 17-producing CD4+ effector T cells develop via a lineage distinct from the T helper type 1 and 2 lineages. Nat Immunol 6, 1123–1132 (2005).

51. Yang, X.O. et al.. Molecular antagonism and plasticity of regulatory and inflammatory T cell programs. Immunity 29, 44–56 (2008).

52. D’Elios, M.M. et al.. Predominant Th1 cell infiltration in acute rejection episodes of human kidney grafts. Kidney Int 51, 1876–1884 (1997).

53. Obata, F. et al.. Contribution of CD4+ and CD8+ T cells and interferon-gamma to the progress of chronic rejection of kidney allografts: the Th1 response mediates both acute and chronic rejection. Transpl Immunol 14, 21–25 (2005).

54. Sadeghi, M. et al.. Pre-transplant Th1 and post-transplant Th2 cytokine patterns are associated with early acute rejection in renal transplant recipients. Clin Transplant 17, 151–157 (2003).

55. Scozzi, D. et al.. The Role of Neutrophils in Transplanted Organs. Am J Transplant 17, 328–335 (2017).

56. Sawant, K.V. et al.. Chemokine CXCL1 mediated neutrophil recruitment: Role of glycosaminoglycan interactions. Sci Rep 6, 33123 (2016).

57. Disteldorf, E.M. et al.. CXCL5 drives neutrophil recruitment in TH17-mediated GN. J Am Soc Nephrol 26, 55–66 (2015).

58. Segerer, S., Nelson, P.J. & Schlöndorff, D. Chemokines, chemokine receptors, and renal disease: from basic science to pathophysiologic and therapeutic studies. J Am Soc Nephrol 11, 152–176 (2000).

59. Paust, H.J. et al.. Chemokines play a critical role in the cross-regulation of Th1 and Th17 immune responses in murine crescentic glomerulonephritis. Kidney Int 82, 72–83 (2012).

60. Stroo, I. et al.. Chemokine expression in renal ischemia/reperfusion injury is most profound during the reparative phase. Int Immunol 22, 433–442 (2010).

61. Lu, G. et al.. CCL20 secreted from IgA1-stimulated human mesangial cells recruits inflammatory Th17 cells in IgA nephropathy. PLoS One 12, e0178352 (2017).

62. Hummel, M. & Abecassis, M.M. A model for reactivation of CMV from latency. J Clin Virol 25 Suppl 2, S123-136 (2002).

63. Hargett, D. & Shenk, T.E. Experimental human cytomegalovirus latency in CD14+ monocytes. Proc Natl Acad Sci U S A 107, 20039–20044 (2010).

64. O’Connor, C.M. & Murphy, E.A. A myeloid progenitor cell line capable of supporting human cytomegalovirus latency and reactivation, resulting in infectious progeny. J Virol 86, 9854–9865 (2012).

65. Reeves, M.B. & Compton, T. Inhibition of inflammatory interleukin-6 activity via extracellular signal-regulated kinase-mitogen-activated protein kinase signaling antagonizes human cytomegalovirus reactivation from dendritic cells. J Virol 85, 12750–12758 (2011).

66. Alcendor, D.J., Charest, A.M., Zhu, W.Q., Vigil, H.E. & Knobel, S.M. Infection and upregulation of proinflammatory cytokines in human brain vascular pericytes by human cytomegalovirus. J Neuroinflammation 9, 95 (2012).

67. Bolovan-Fritts, C.A., Trout, R.N. & Spector, S.A. Human cytomegalovirus-specific CD4+-T-cell cytokine response induces fractalkine in endothelial cells. J Virol 78, 13173–13181 (2004).

68. Shimamura, M., Murphy-Ullrich, J.E. & Britt, W.J. Human cytomegalovirus induces TGF-beta1 activation in renal tubular epithelial cells after epithelial-to-mesenchymal transition. PLoS Pathog 6, e1001170 (2010).

69. Grundy, J.E., Lawson, K.M., MacCormac, L.P., Fletcher, J.M. & Yong, K.L. Cytomegalovirus-infected endothelial cells recruit neutrophils by the secretion of C-X-C chemokines and transmit virus by direct neutrophil-endothelial cell contact and during neutrophil transendothelial migration. J Infect Dis 177, 1465–1474 (1998).

70. Nordøy, I. et al.. Chemokines and soluble adhesion molecules in renal transplant recipients with cytomegalovirus infection. Clin Exp Immunol 120, 333–337 (2000).

71. Annunziato, F. et al.. Phenotypic and functional features of human Th17 cells. J Exp Med 204, 1849–1861 (2007).

72. Duhen, T. & Campbell, D. Human CCR6+CXCR3+ “Th1/17” cells are polyfunctional and can develop from Th17 cells under the influence of IL-1β and IL-12 (P4122). The Journal of Immunology 190, 191.197-191.197 (2013).

73. Harbour, S.N., Maynard, C.L., Zindl, C.L., Schoeb, T.R. & Weaver, C.T. Th17 cells give rise to Th1 cells that are required for the pathogenesis of colitis. Proc Natl Acad Sci U S A 112, 7061–7066 (2015).

74. Zielinski, C.E. et al.. Pathogen-induced human TH17 cells produce IFN-gamma or IL-10 and are regulated by IL-1beta. Nature 484, 514–518 (2012).

75. Tsanaktsi, A., Solomou, E.E. & Liossis, S.C. Th1/17 cells, a subset of Th17 cells, are expanded in patients with active systemic lupus erythematosus. Clin Immunol 195, 101–106 (2018).

76. Wilson, N.J. et al.. Development, cytokine profile and function of human interleukin 17- producing helper T cells. Nat Immunol 8, 950–957 (2007).

77. Amadi-Obi, A. et al.. TH17 cells contribute to uveitis and scleritis and are expanded by IL-2 and inhibited by IL-27/STAT1. Nat Med 13, 711–718 (2007).

78. Santarlasci, V. et al.. TGF-beta indirectly favors the development of human Th17 cells by inhibiting Th1 cells. Eur J Immunol 39, 207–215 (2009).

79. Veldhoen, M., Hocking, R.J., Atkins, C.J., Locksley, R.M. & Stockinger, B. TGFbeta in the context of an inflammatory cytokine milieu supports de novo differentiation of IL-17-producing T cells. Immunity 24, 179–189 (2006).

80. Kortylewski, M. et al.. Regulation of the IL-23 and IL-12 balance by Stat3 signaling in the tumor microenvironment. Cancer Cell 15, 114–123 (2009).

81. Khader, S.A. et al.. IL-23 and IL-17 in the establishment of protective pulmonary CD4+ T cell responses after vaccination and during Mycobacterium tuberculosis challenge. Nat Immunol 8, 369–377 (2007).

82. Haines, C.J. et al.. Autoimmune memory T helper 17 cell function and expansion are dependent on interleukin-23. Cell Rep 3, 1378–1388 (2013).

83. Nazari, B. et al.. Comparison of the Th1, IFN-γ secreting cells and FoxP3 expression between patients with stable graft function and acute rejection post kidney transplantation. Iran J Allergy Asthma Immunol 12, 262–268 (2013).

84. Dobin, A. et al.. STAR: ultrafast universal RNA-seq aligner. Bioinformatics 29, 15–21 (2013).

85. Love, M.I., Huber, W. & Anders, S. Moderated estimation of fold change and dispersion for RNA-seq data with DESeq2. Genome Biol 15, 550 (2014).

86. Loupy, A. et al.. The Banff 2019 Kidney Meeting Report (I): Updates on and clarification of criteria for T cell- and antibody-mediated rejection. Am J Transplant 20, 2318–2331 (2020).

87. Roufosse, C. et al.. A 2018 Reference Guide to the Banff Classification of Renal Allograft Pathology. Transplantation 102, 1795–1814 (2018).

88. Dhital, R. et al.. Flow cytometric quantitation of T-cell subsets in peripheral blood of renal allograft recipients. Indian Journal of Transplantation 10 (2016).

89. Dhital, R. et al.. OR19 Quantification of peripheral B-cell subsets in acute allograft rejection in recipients with renal transplantation. Human Immunology 77 (2016).

90. Minz, M., Dhital, R., Jha, V., Minz, R.W., Sharma, A. mRNA expression of BAFF and APRIL receptors increases in acute rejection in kidney transplant recipients. Am J Transplant 16 (2016).

91. Mitra, S. et al.. A molecular marker of disease activity in autoimmune liver diseases with histopathological correlation; FoXp3/RORgammat ratio. APMIS 123, 935–944 (2015).

92. Minz, R.W. et al.. Cytomegalovirus Infection in Postrenal Transplant Recipients: 18 Years’ Experience From a Tertiary Referral Center. Transplant Proc 52, 3173–3178 (2020).

93. Edgar, R., Domrachev, M. & Lash, A.E. Gene Expression Omnibus: NCBI gene expression and hybridization array data repository. Nucleic Acids Res 30, 207–210 (2002).

